# Study design and the sampling of deleterious rare variants in biobank-scale datasets

**DOI:** 10.1101/2024.12.02.626424

**Authors:** Margaret C. Steiner, Daniel P. Rice, Arjun Biddanda, Mariadaria K. Ianni-Ravn, Christian Porras, John Novembre

**Author notes:** **Corresponding author:** John Novembre CLSC 421, 920 E 58th St, Chicago, IL 60637. These authors contributed equally to the work (Margaret C. Steiner, Daniel P. Rice).

## Abstract

One key component of study design in population genetics is the “geographic breadth” of a sample (i.e., how broad a region across which individuals are sampled). How the geographic breadth of a sample impacts observations of rare, deleterious variants is unclear, even though such variants are of particular interest for biomedical and evolutionary applications. Here, in order to gain insight into the effects of sample design on ascertained genetic variants, we formulate a stochastic model of dispersal, genetic drift, selection, mutation, and geographically concentrated sampling. We use this model to understand the effects of the geographic breadth of sampling effort on the discovery of negatively selected variants. We find that samples which are more geographically broad will discover a greater number variants as compared geographically narrow samples (an effect we label “discovery”); though the variants will be detected at lower average frequency than in narrow samples (e.g. as singletons, an effect we label “dilution”). Importantly, these effects are amplified for larger sample sizes and moderated by the magnitude of fitness effects. We validate these results using both population genetic simulations and empirical analyses in the UK Biobank. Our results are particularly important in two contexts: the association of large-effect rare variants with particular phenotypes and the inference of negative selection from allele frequency data. Overall, our findings emphasize the importance of considering geographic breadth when designing and carrying out genetic studies, especially at biobank scale.

**Significance:** As genetic studies grow, researchers are increasingly seeking to identify rare genetic variants with large impacts on traits. In this paper, we combine theoretical methods and data analysis to show how differences in sampling with respect to geographic location can influence the number and frequency of genetic variants that are found. Our results suggest that geographically broad samples will include more distinct genetic variants, though each variant will be found at a lower frequency, as compared to geographically narrow samples. Our results can help researchers to consider the implications of study design on expected results when constructing new genetic samples.

## Introduction

In recent decades, the size of genetic sequencing cohorts has grown exponentially. Nowhere is this more evident than in human genetics, where the launch of biobanks has transformed the paradigm of data analysis such that sample sizes in the hundreds of thousands are increasingly commonplace (Gallagher et al., 2024). Yet, the largest and most commonly utilized biobank-scale genomics datasets are heavily biased towards individuals of European ancestries (Bustamante et al., 2011; Popejoy and Fullerton, 2016; Bycroft et al., 2018; Karczewski et al., 2020), leading to known issues in scientific discovery and ethical applications of precision medicine (Martin et al., 2019; Dolan et al., 2023). As a response to this, new biobanks have been launched with specific purposes to diversify available genomics data (Sohail et al., 2023; All of Us Research Program Genomics Investigators, 2024; Verma et al., 2024; Elfatih et al., 2024). Consequently, not only is the size of human genetics data continuing to increase, but the geographic and genetic spaces from which individuals are sampled is growing dramatically.

This trend in the field has clear benefits for improving equity in human genetics research and the transferability of results across diverse populations (Sirugo et al., 2019; Durvasula and Lohmueller, 2021; Ding et al., 2022). What has yet to be addressed is how this change in study design will affect the results of genetic studies at the level of discovered variants. Motivated as such, we ask: as the *geographic breadth* of a genetic study increases, how should one expect the number and frequency of discovered variants to change? That is, how is the site frequency spectrum (SFS) of observed variants affected by the geographic breadth of a sample? The answer to this question has significant implications for studies in human genetics and more broadly.

For understanding the genetic basis of traits, this question is of interest because sample design likely impacts the discovery of genetic associations to phenotypes. A key focus of biomedical applications is discovering variants that have large effects on disease susceptibility, as such variants may provide the most biological insight on the etiology of disease and in turn potential therapeutic paths (Szustakowski et al., 2021; Ghoussaini et al., 2023). From evolutionary principles, one expects large effect genetic changes most often to be kept at very low population frequencies by the action of natural selection (either due to simple negative selection or via underdominance induced by stabilizing selection; Sella and Barton, 2019). Indeed, rare, deleterious variants have been shown to be enriched in genomic regions of functional interest such as drug target regions (Weiner et al., 2023), have yielded numerous associations with phenotypic outcomes (Backman et al., 2021; Sun et al., 2022), and are argued to be a key component of unexplained heritability in human traits (Wainschtein et al., 2022). How the geographic breadth of sampling impacts the discovery of these rare, deleterious variants is unknown, yet crucial to the design of studies which aim to characterize such variants.

Understanding how sample design affects the observed SFS of deleterious variants is also important to evolutionary geneticists. In evolutionary genetics, a persistent goal has been to characterize the distribution of fitness effects (DFE) – i.e., the probability with which newly arising mutations are deleterious, advantageous, or selectively neutral – using allele frequency data (Williamson et al., 2005; Boyko et al., 2008; Gutenkunst et al., 2009; Kim et al., 2017), in part because of its implications for genome evolution, mutational load, and conservation efforts (Robinson et al., 2023). Population-genetic-based inferences regarding the DFE depend on the measurement of the numbers and frequencies of deleterious variants (equivalently, the SFS). Thus, whether and how the geographic breadth of sampling impacts the observed SFS of deleterious variants is also important for evolutionary geneticists to understand, in order to avoid biases in SFS-based inferences of fitness effects.

Previous studies, motivated by understanding the consequences of spatial structure and sampling on the inference of demographic history, have investigated the effect of sample design on *neutral* variation (Wakeley, 1999; Ptak and Przeworski, 2002; Arunyawat et al., 2007; Städler et al., 2009; Quéméré et al., 2010; St Onge et al., 2012; Mazet et al., 2015; Battey et al., 2020; Gloss et al., 2022). These studies emphasize how in most cases, geographically concentrated (or “narrow”) sampling in spatial populations leads to a shift in the neutral SFS with a decrease in observed singletons and enrichment of intermediate and high frequency alleles (i.e. negative Tajima’s D; Tajima, 1989). These previous studies do not consider sample sizes on the scale of modern human biobank cohorts, which reach tens to hundreds of thousands of individuals, nor do they address the extent to which distortions of the SFS are amplified or diminished for rare, deleterious variants.

Here, with a focus on the discovery of rare, deleterious variants, we develop and analyze a novel theoretical model for the sample SFS in a spatially structured population. The model considers the distribution of carriers of deleterious alleles in continuous geographic space – accounting for dispersal, genetic drift, selection, mutation, and sampling simultaneously – and we derive results that allow the rapid computation of the expected sample SFS across a large range of parameter values.

As important background, we note that in the panmictic case, allele frequencies for deleterious variants are well known to follow a two-parameter distribution, such that the probability *g*(*x*) that an allele under negative selection appears at frequency *x* follows (Wright, 1937; Crow and Kimura, 1970):

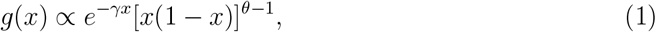

where *γ* is the population-scaled selection coefficient (*γ* = 4*N*_*e*_*s* with *N*_*e*_ being the effective population size and *s* is the strength of negative selection acting on heterozygotes, *s* ≥ 0) and *θ* is the population-scaled mutation rate (*θ* = 4*N*_*e*_*µ* with mutation rate *µ* per site per generation). For variants under negative selection (*γ* > 0), the exponential term (*e*^−*γx*^) induces a reduction in the abundance of observed alleles as a function of the allele frequency *x*. This mirrors the intuition that alleles under negative selection are less likely to reach higher frequencies.

As we will show, when considering spatially-restricted dispersal and geographically concentrated sampling, allele frequencies still follow a two-parameter distribution with scaled selection (*γ*_*E*_) and mutation (*θ*_*E*_) terms. However, these terms are now dependent on the spatial scale of the sampling effort and the offspring dispersal scale, in addition to the usual mutation, selection, and population size parameters. The resulting distributions show that the geographic breadth of a sample has strong effects on the SFS as well as downstream summary statistics, and we assess how these effects change with increasing sample size and selection strength.

We validate our theoretical results using simulations that share our modeling assumptions as well as in a more realistic, individual-based spatial model (Haller and Messer, 2019; Battey et al., 2020). However, as continuous-space simulations can be computationally intensive, our development of novel theoretical approximations allows us to efficiently gain insights across a wide parameter range.

To address the effects of geographically concentrated sampling empirically, we also conduct in silico re-sampling experiments using the UK Biobank exome sequencing dataset to measure the impact of sampling at different scales – in either geographic space or a low-dimensional genetic space (e.g., a PCA space). The results broadly confirm our theoretical predictions and yield insights on how sampling design impacts the number of human genetic variants discovered and their frequencies.

## Methods

### Population genetic model

We model how carriers of a rare variant are born, move in space, reproduce, and die in a two-dimensional continuous geographic habitat of size *L × L*. In our model (Fig. 1), carriers are generated by *de novo* mutation according to a Poisson point process with rate *ρ*_*N*_ *µ*, where *ρ*_*N*_ is the population density and *µ* is the per-generation mutation rate. We note that, in our model, *ρ*_*N*_ and *L* are constants, which implies an assumption of constant population size. Each *de novo* carrier appears at a random location in the habitat and migrates according to a homogeneous, isotropic diffusion process (i.e., the dispersal process is the same across the habitat and movement is uniform in all directions). The root mean squared distance that one carrier moves per generation is denoted by *σ*. Similar to De and Durrett (2007), the habitat has periodic boundary conditions (i.e., the habitat is a two-dimensional torus, and so there will be no boundary effects). While our model is defined over continuous geographic space – which has some advantages in that it more closely resembles realistic geography and allows us to utilize a simple model of migration – we note that it is possible to develop the analysis in terms of a demic model, such as a stepping stone model. Such models approximate continuous space when the grid of demes is dense (Petkova et al., 2016; Al-Asadi et al., 2019; Marcus et al., 2020).

**Figure 1:**
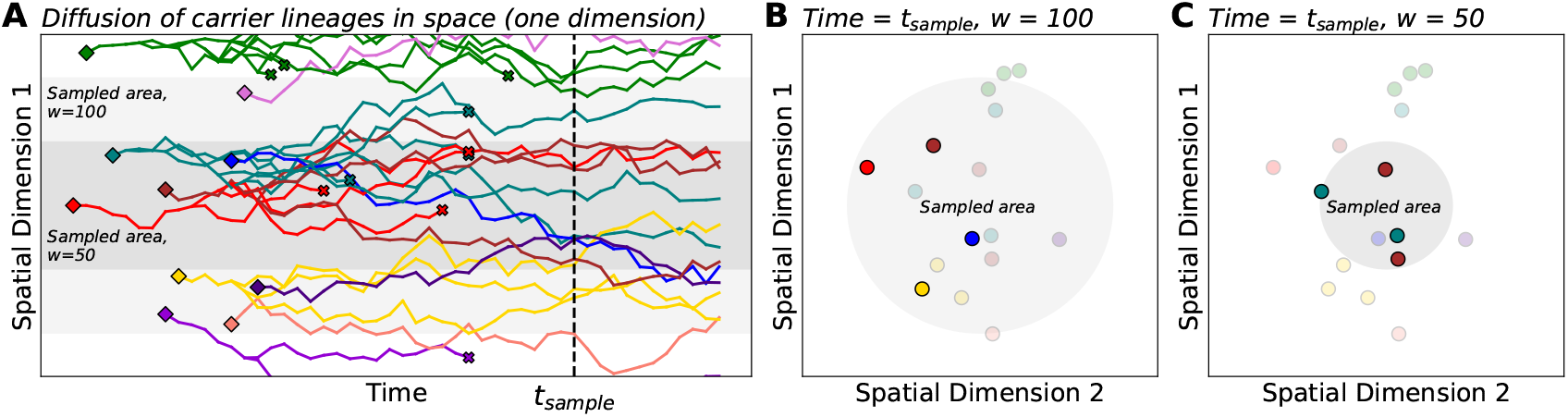
Illustration of a spatial branching process model with sampling. (**A**) As time progresses, carrier lineages move in space (diffusion), branch into sub-lineages (reproduction), and die. Diamonds and x’s denote lineage birth and death, respectively. For simplicity, we show only one spatial dimension on the vertical axis. Shaded areas represent sampled areas with widths *w* = 50 and *w* = 100. (**B, C**) Here, we visualize the locations of carriers from panel **A** at a particular time indicated as *t*_*sample*_. Within the sampled area, rare variant carriers can be discovered and included in the sample (opaque points). In this example, sampling from the broader area (*w* = 100) results in a greater number of distinct mutations being observed, all as singletons. The narrow sample (*w* = 50) discovers two distinct mutations with each as a pair of doubletons. This toy example illustrates the potential effects of sampling breadth on entries of the sample SFS (here, the counts of singletons vs. doubletons).

Within the habitat, we model reproduction and death as a continuous time branching process, a type of stochastic process which has frequently been applied in theoretical population genetics for rare variants (see, for instance, Ewens, 1968; Tavaré, 1984; Peter and Slatkin, 2015; Etheridge et al., 2017). During their lifetime, carriers reproduce to form offspring carriers with rate 1 − *s* and die at a rate 1, where *s* denotes the fitness cost to carriers of the mutation (note that in our construction, a larger *positive* value of *s* indicates stronger *negative* selection). While we model negative selection on individual variants, these dynamics are similar to those of newly arising variants which affect complex traits under stabilizing selection which experience a form of underdominance (Sella and Barton, 2019). The use of a branching process model implies that carriers evolve independently of one another and of wild-type individuals (similar to Haldane, 1927; Slatkin and Rannala, 1997; Novembre and Slatkin, 2009). In the context of continuous space, this approximations will hold best when every mutation is locally rare (more precisely, the number of carriers in a region of the habitat with radius *σ* is small compared to the neighborhood size, 4*πσ*^2^*ρ*; Wright, 1946).

### Modeling geographically concentrated sampling

The spatial model in the previous section describes the process by which rare variants arise and disperse in geographic space. Our next step is to define how sampling of this spatial population occurs. To this end, we model the probability that an individual at a particular position within the habitat is included in the sample. We posit a sampling “center” and assume that the probability of being sampled is determined by an individual’s distance from that center using a particular distribution (the “sampling kernel”; Fig. S1). The standard deviation of the sampling kernel, which we denote by *w*, determines the breadth of sampling effort – or “sampling breadth” – i.e. the extent to which sampling effort is distributed across the habitat. On one extreme, for *w* ≳ *L*, the sampling process converges to “uniform” sampling in which all individuals have an equal probability of being sampled, regardless of spatial position. In the other extreme, as *w* becomes small, the sampling kernel approaches “point sampling” in which all sampled individuals are located at the same position. In between these two limiting cases, the value of *w* will determine how spatially “broad” (larger *w*) or “narrow” (smaller *w*) a sample will be.

In our implementation, the sampling kernel has the form of a Gaussian distribution, though we note that our methods are generalizable to other sampling kernels (see Supplemental Information 1.1.2). We employ the Gaussian sampling model to approximate the sampling processes used in constructing real genetic samples, such as sampling centered at field stations for ecological genetics or in biomedical centers for human genetics. We also invoke periodic boundary conditions for mathematical convenience (i.e., there are no “edge effects” in our model). This construction is most appropriate when the habitat size, *L*, is sufficiently large compared to the sampling breadth, *w*, such that we can ignore behavior as the sampling kernel hits the edge of the habitat (Our simulations will show that in cases where *w* approaches the scale of *L* , the results converge to those of uniform random sampling).

In order to solve for the SFS, we obtain moments of an allele frequency distribution at equilibrium that incorporates the spatially-weighted sampling design, but assumes infinite sample sizes. We do so by casting our model of a spatial branching process with spatially-weighted sampling as an example of a *superprocess* (see Etheridge, 2000; A. M. Etheridge, 2004; Etheridge et al., 2017). The moments allow us to approximate the full spatially-weighted allele frequency distribution at equilibrium, which we then use to calculate the expected observed SFS for a finite sample of size *n*. Finally, we use the finite sample observed SFS to derive expectations population genetic summary statistics. In essence, we first consider the effects of geographically concentrated sampling on the SFS without invoking finite sampling, and then, we compute the expected observed SFS for a finite sample of size *n*.

### Population genetic simulations

We validate our theoretical results with two sets of simulations. First, we simulate a spatial branching process in a two-dimensional continuous habitat and sample according to a Gaussian sampling density, as our theory assumes. These simulations are close to our theory in that they make the same rare-allele approximation. Their role is to check the analytical approximations we make in the course of deriving the SFS.

In addition to the branching process simulations, we implement out-of-model, forward-time, individual-based population genetic simulations in SLiM (Haller and Messer, 2019) using identical conditions to Battey et al. (2020) except that all variants are deleterious with some selection coefficient. For each simulation run, we sample individuals using Gaussian sampling kernels with varying standard deviation and calculate the sample SFS. In contrast to our other simulations, the SLiM model contains multiple stages of the life cycle, models diploid genomes, and – crucially – does not assume variants are rare and independently evolving.

We refer the reader to the Supplemental Information for additional details on simulation methods. All simulation code and associated scripts are available at: https://github.com/NovembreLab/spatial rare alleles.

### Analysis of whole exome sequencing data from the UK Biobank

We perform re-sampling experiments using the whole exome sequencing (WES) dataset (*n* = 469, 835) in the UK Biobank (UKB; Backman et al., 2021) in order to assess the predicted effects of sampling breadth on sample allele frequencies and associated summary statistics. We first compute the top 20 PCs using genotyping array data via PLINK (v2.00a3.1LM), including only individuals which met quality control and relatedness thresholds used in Bycroft et al. (2018). We then take two approaches to our empirical investigations: geographic sampling by birthplace and PCA-based sampling.

For the geographic sampling approach, we utilize the birthplace coordinates provided by UKB and subset to individuals both born within the UK and having similar genetic ancestry (specifically, individuals within 0.0001 of the centroid in the normalized PC1-PC2 space; applying both filters results in *n* = 231, 073 individuals). We then use a sampling importance resampling (SIR) method to construct *n* = 10, 000 samples such that the distribution of birthplace coordinates is Gaussian with centers centered at each of three geographic points with standard deviation 50km, 100km, and 150km, as well as a uniform sample (see Supplemental Information 1.3.1 for details on the sampling algorithm). For sampling in PCA space, we center the distribution of sample (PC1, PC2) coordinates at the centroid of PC1-PC2 space and construct *n* = 10, 000 Gaussian samples with standard deviations 0.0015, 0.0025, and 0.005, as well as a uniform sample. In both cases, we repeat the sampling procedure ten times for each sampling width and center (for samples in geographic space only).

For each weighted subsample, we compute the site frequency spectrum for LoF variants on chromosome 1 (54,090 variants) as well as equal-sized random subsets of synonymous and missense variants (created using PLINK v1.90b6.26). We then use the computed frequency spectra to calculate summary statistics (number of variant sites, number of singletons, and allele frequency for variant sites only vs. all sites).

## Results

### The finite sample SFS depends on ratios between spatial scales as well as sample size

In our model, a key emergent feature is the distance an allele spreads during the time from the initial mutation to the extinction of all its carriers, which we denote as ℓ_*c*_ (the *characteristic length scale*). As the carrier lineages diffuse at rate *σ* and the time-scale of allele age is on average 1*/s* generations, the scale ℓ_*c*_ is naturally 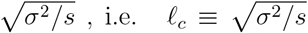. Intuitively, carriers which spread more quickly (large *σ*) can move farther distances during the lifespan of the allele (ℓ_*c*_ is large). Conversely, alleles which are under stronger negative selection (large *s*), die more quickly and thus carriers move shorter distances (ℓ_*c*_ is small). As we see later on in our results, how the spatial scale of the sampling effort (*w*) compares to the spatial spread of the allele (ℓ_*c*_) will be an important factor in the behavior of the SFS.

In order to derive the form of the SFS for a finite sample of size *n* with sampling effort breadth *w*, we first consider the distribution of allele frequencies across the entire spatially extended population with a weighting on each position provided by the sampling kernel. In a panmictic population, the population SFS of rare deleterious alleles approximately follows a gamma density (by ignoring the *x* → 1 tail of Eq. 1). We show analytically that this approximation also holds for spatially uniform samples under our model (Supplemental Information 1.1.6) and confirm via simulations that allele frequencies of spatially concentrated samples are also well-approximated by a gamma distribution (Fig. S2-S3). Intuitively, the gamma distribution captures two important effects: power-law behavior at low frequencies due to mutation-drift balance, and an exponentially decaying tail at high frequencies due to selection.

We analyze our model in order to obtain the two parameters of this gamma distribution, which we refer to as the effective mutation supply, *θ*_*E*_, and the effective selection intensity, *γ*_*E*_ (see Supplemental Information 1.1.4). Then, we derive an expression for the expected SFS of a finite sample with size *n* in terms of these parameters. First, let the random variable *K* denote the count of derived alleles at a single site in a sample of size *n*. Combining our results regarding the allele frequency distribution with a Poisson approximation to the binomial sampling process, *K* follows a Negative Binomial distribution with number of successes *θ*_*E*_ and success probability *γ*_*E*_*/*(*γ*_*E*_ + *n*):

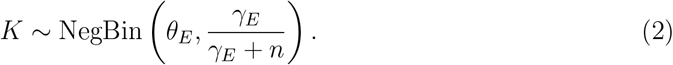

The *k*-th entry of the normalized sample SFS is then given by 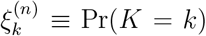. In the limit that *θ*_*E*_ is small, this becomes approximately:

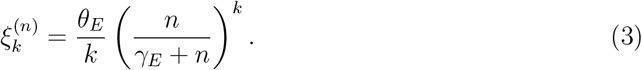

In the case where sampling is approximately spatially uniform (*w* ≈ *L* or larger), we find that (similarly to Eq. 1), both terms take the form of population-scaled parameters: *θ*_*E*_ = *Nµ* and *γ*_*E*_ = *Ns*, respectively, for *N* the total population size. As such, in the spatially uniform sampling case, our results are equivalent to that of the panmictic case.

In the case of spatially concentrated sampling (*w* << *L*), we show (see Supplemental Information 1.1.4) that the effective mutation and selection terms are instead given by:

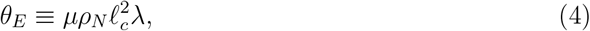

and

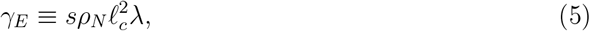

Where 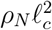 is akin to a population size and λ is term we denote as the *sampling effect scalar* that is a function of *w/*ℓ_*c*_:

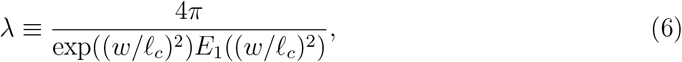

where *E*_1_(*x*) is the exponential integral function. The λ parameter increases as *w* increases (Fig. 2 A). As a result, both effective parameters also increase monotonically with *w* ,and eventually converge to their values in the uniform sampling limit (*Nµ* and *Ns*, respectively; Fig. 2 C,D). To understand this result, one can think of the term 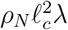 as approximating the size of the population effectively being sampled (which converges to *N* as sampling converges to the uniform case). Comparing the finite sample SFS for *w* << *L* to Eq. 1, a key difference is that the effective selection parameter (*γ*_*E*_) is moderated by *w* via its impact on ℓ_*c*_ and λ.

**Figure 2:**
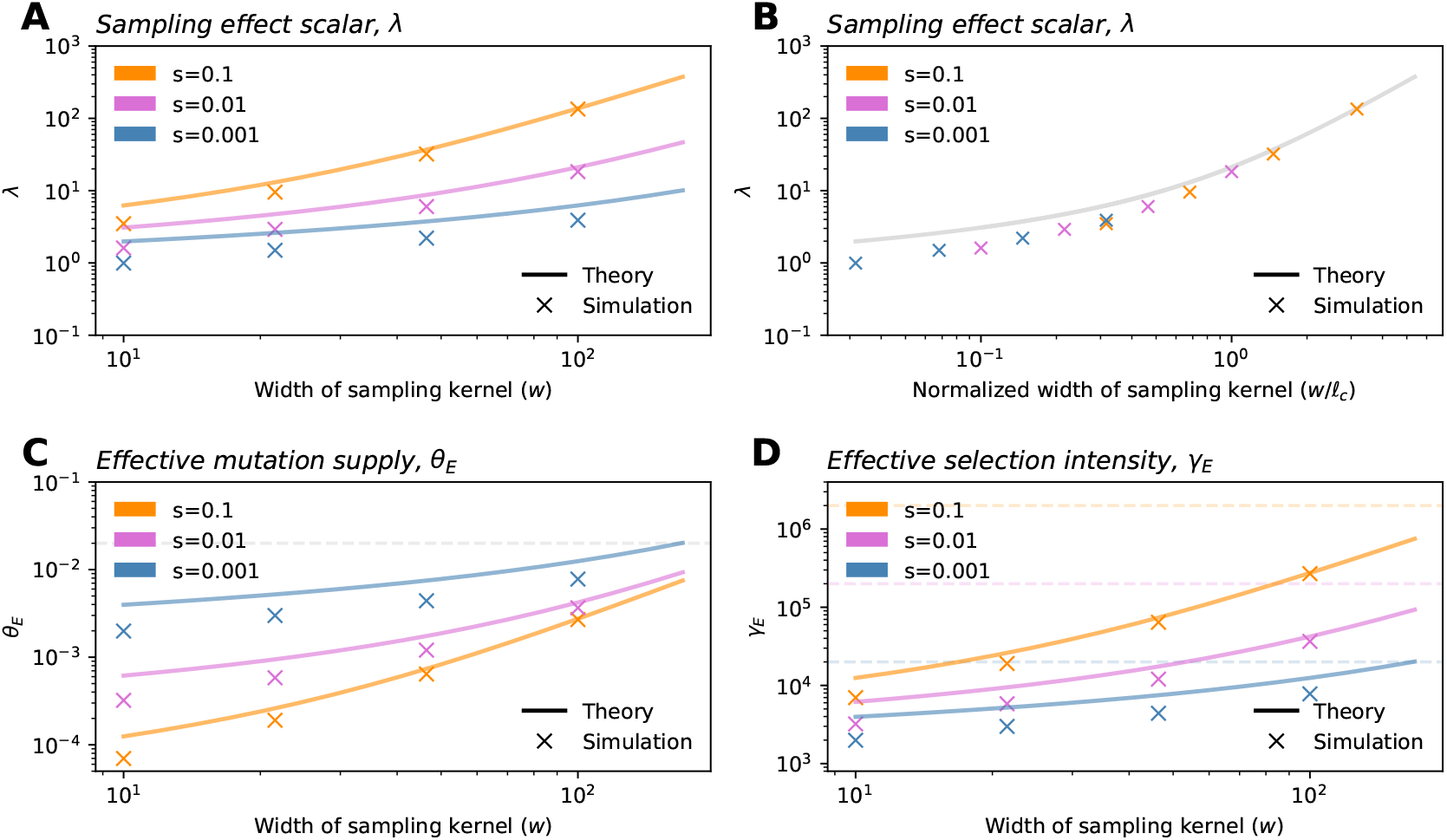
The sampling effect scalar and effective parameters of the SFS. (**A**) As the breadth of the sampling kernel increases, the sampling effect scalar λ also increases. When plotted as a function of *w/*ℓ_*c*_, the relationship is identical across selection intensities (**B**). (**C**-**D**) Both the effective mutation supply, *θ*_*E*_, and the effective selection intensity, *γ*_*E*_, depend on the selection coefficient (via ℓ_*c*_) and the breadth of the sampling kernel. Dashed lines show values of *θ*_*E*_ and *γ*_*E*_ for the uniform case in panels **C** and **D**, respectively. Other parameters are: *σ* = 10, *ρ* = 20, and *µ* = 1*e* − 9. All simulations shown are from the Gillespie algorithm run with a habitat size of *L* = 1, 000.

To summarize, one ratio between length scales – the ratio between *w* and *L* – determines the regime in which the finite sample SFS lies. For *w* ≳ *L*, sampling is approximately uniform and the SFS has approximately the same form as in the panmictic case. For *w* << *L*, the form of the SFS is instead dependent on the value of the sampling effect scalar, λ, which is a function of a second ratio: *w/*ℓ_*c*_. We note that these ratios between spatial terms are dimensionless (appropriately, if one switches the units of space from miles to kilometers, the SFS should not change).

### Selection and sampling induce a trade-off between discovery and dilution

Having derived an expression for the sample SFS, we now consider its behavior with respect to the sampling width (Fig. 3 A, B). We find broader sampling effort (larger *w*) induces an upward shift in the intercept of the SFS on the vertical axis, and this effect is more apparent in larger samples. We also observe a decrease in the relative frequency of intermediate-frequency variants for broader samples. Additionally, as sampling effort broadens (*w* increases), the SFS converges to the result under uniform sampling, as expected.

**Figure 3:**
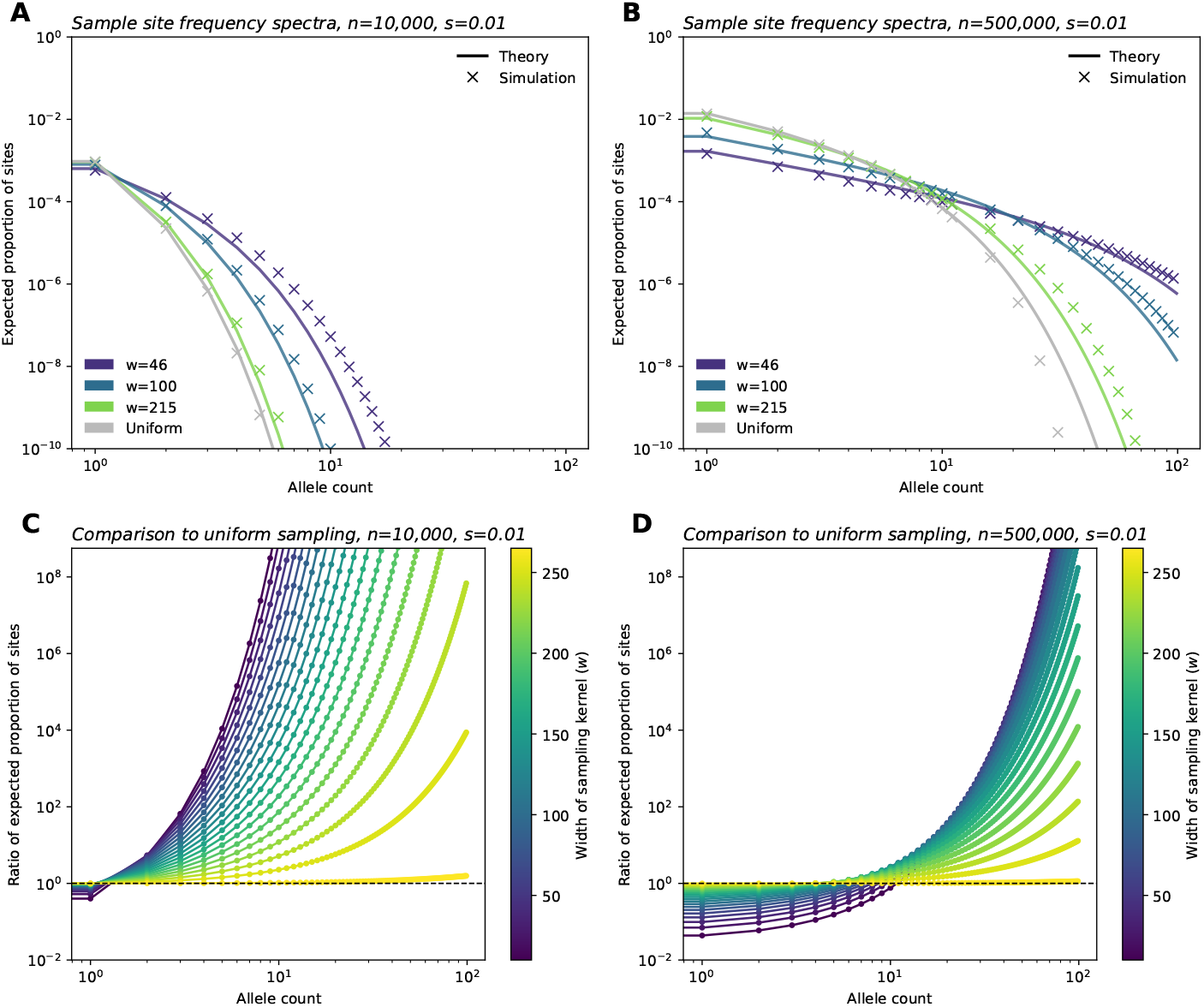
Site frequency spectra for a range of sampling widths. (**A**,**B**) Sample site frequency spectra for *n* = 10, 000 and *n* = 500, 000 sample sizes, respectively, shown for three sampling widths (wrapped Gaussian sampling) and uniform sampling. (**C**,**D**) Ratio between frequency spectra elements for a range of *w* values (Gaussian sampling), relative to those of the SFS under uniform sampling, for parameter regimes shown in **A** and **B**. For all panels, *σ* = 10, *ρ* = 20, *µ* = 1*e* − 9 and *s* = 0.01. All simulations shown were run with a habitat size of *L* = 1, 000.

Another way to understand the impact of sample breadth on the SFS is to recall that as *w* increases (i.e., sampling becomes more broad), both *θ*_*E*_ and *γ*_*E*_ increase (Fig. 2). This results in what we term a “discovery” effect and a “dilution” effect. As the geographic breadth of a sample increases, the number of potential localized mutations one can find grows, and this is reflected in the increase in mutation supply (*θ*_*E*_), as well as resulting increase in the expected number of variants discovered (the *discovery effect* ). At the same time, for a broader sample, each sampled deleterious variant is found at “diluted” frequencies because sampling broadly inadvertently captures many non-carriers (given that deleterious variants are usually spatially clustered). This is reflected in the fast rate of decay with *k* owing to the larger *γ*_*E*_ term (*dilution effect* ). Conversely, geographically narrow samples will capture fewer variants, but they are “concentrated” in the sample, meaning that they are observed at higher sampled allele frequencies on average than they would be found otherwise in a random sample of the population.

We visualize these results by comparing each entry of the SFS of a sample with breadth *w* to the SFS in the uniform limit (Fig. 3 C,D). Below a threshold value of allele count, we expect to fewer variants in the narrower samples. Above the threshold, we expect to see more variants at these counts in the narrower sample. The location of this threshold is dependent on the sample size, *n*: for smaller samples the threshold is low, perhaps only affecting singletons; for larger samples, the threshold is higher, with a larger range of rare allele counts being less often sampled in the narrow relative to the uniform sample. The magnitude of effect in the low allele count range is also much larger for the larger sample.

The changes to the observed SFS with sampling breadth have varying effects on downstream summary statistics (Fig. 4). First, we see that broader samples will have a greater proportion of variant sites and singletons as compared to narrower samples. Secondly, variant sites in broader samples are expected to have lower allele frequency. Together, these two results imply that broader samples will have more variants, but each variant will segregate at lower frequency on average. This result is consistent with intuition following the discovery and dilution effects described previously. Each of these values converges to the expectation under uniform sampling as *w* increases.

**Figure 4:**
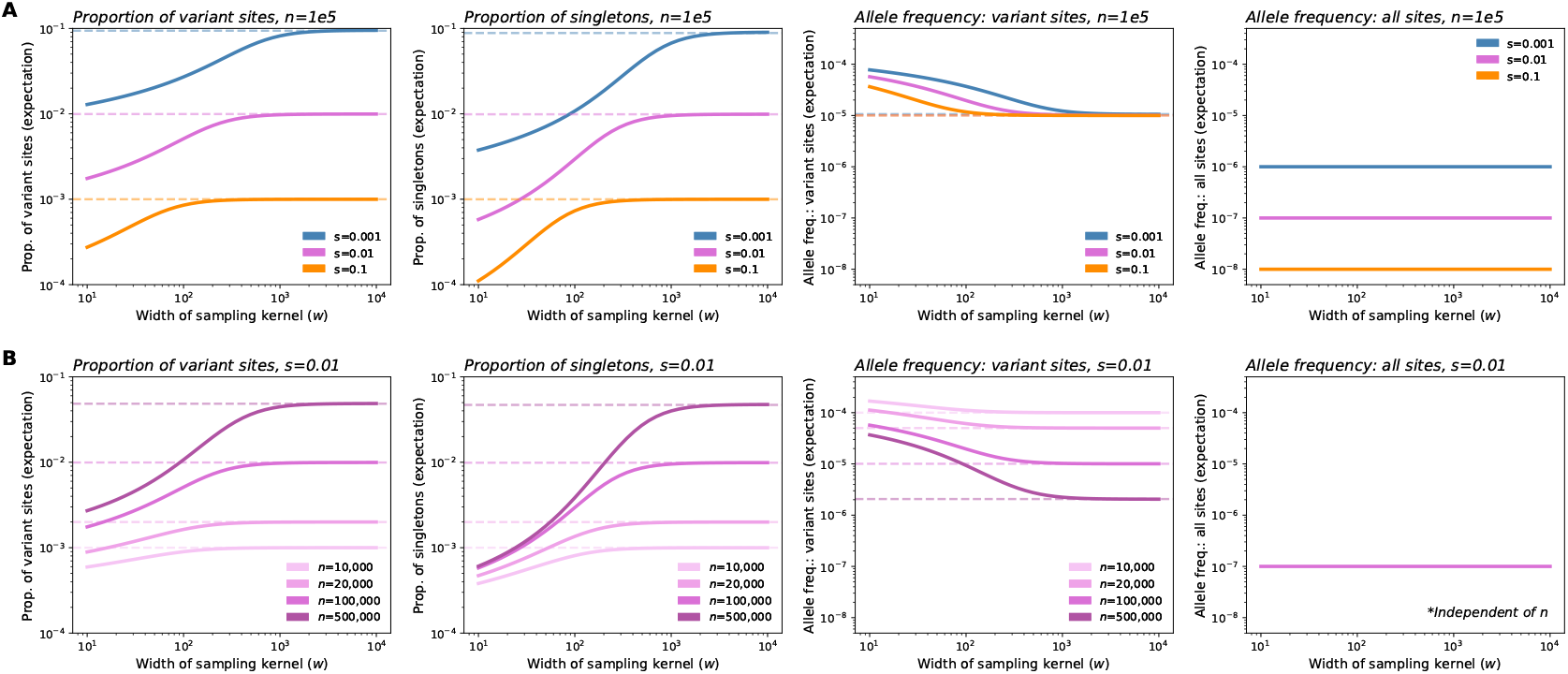
Expected values of summary statistics as a function of the width of the sampling kernel (*w*). (**A**) As the breadth of the sampling kernel increases, our model implies that the expected proportion of variant sites and the expected proportion of singletons will both increase, the expected frequency of variant sites will decrease, while expected frequency across all sites will remain constant. Values for these statistics (excluding expected frequency across all sites) converge to the theoretical expectation under uniform sampling (dashed lines) as *w* increases, with convergence occurring more quickly for stronger selection coefficients. For more deleterious variants, the expected proportions of variant sites and singletons, as well as expected frequency across all sites, are shifted downwards across the range of *w*. (**B**) Fixing *s* and instead varying sample size (*n*), we see that the magnitude of change as *w* increases is higher for larger sample sizes. Expected allele frequency across all sites is independent of sample size. In plots shown, *σ* = 10, *ρ*_*N*_ = 20, and *µ* = 1*e* − 9.

The exact behavior of these statistics with respect to *w* depends on the strength of selection (Fig. 4A). With stronger selection, the observations converge to those expected under uniform sampling more rapidly as *w* is increased. When instead considering the values of expected summary statistics over the ratio *w/*ℓ_*c*_, we see that the rate and magnitude of change are consistent across selection coefficients (Fig. S11). This is a result of the ratio *w/*ℓ_*c*_ being the key length scale in our model (Fig. 2B): for a fixed *w*, stronger selection reduces ℓ_*c*_, because allele carriers are more tightly clustered in space, and as a result, the sample is in effect more broad relative to the spatial dispersion of the carriers. Conversely, with the strength of selection held constant, increasing *w* results in the sample being more broad relative to the spatial dispersion of carriers.

Holding selection constant, we also see that the magnitude of effect as *w* increases becomes larger as *n* increases (Fig. 4B). These effects are quite large, spanning several orders of magnitude. This result is in line with the observation from Fig. 3D that the discovery effect applies to a larger range of allele counts.

However, other quantities of interest are not sensitive to the sampling breadth, including the expected allele frequency of all sites, expected heterozygosity, and expective cumulative MAF (Fig. 4, Fig. S10). In particular, under our model, the expected allele frequency in the sample is equal to *µ/s*, and is independent of sampling strategy, sampling breadth, and sample size. This indicates that the discovery and dilution effects effectively cancel each other out, such that the average frequency (and in turn the expected heterozygosity and the cumulative minor allele frequency of rare variants at a locus) remain the same regardless of sampling (Fig. S10). Expected heterozygosity and cumulative MAF, in turn, are approximately proportional to the average allele frequency (see Supplemental Information 1.1.5).

### Validation of theory using in- and out-of-model simulations

To validate our theoretical results, we performed two sets of population genetic simulations. First, we ran branching process simulations which correspond directly to our model. Inspecting the results, we see the simulations and theoretical computations align well for key outputs: the first two moments of the allele frequency (Fig. S4), the SFS (Fig. 3, Fig. S2-S3), as well as for λ, *θ*_*E*_, and *γ*_*E*_ (Fig. 2).

As a stronger test of the theory, we performed simulations in SLiM (Haller and Messer, 2019) using a modified version of the model in Battey et al. (2020). These simulations are individual-based and do not make a number of the simplifying approximations used in our theoretical analysis. Figure 5 shows the alignment between the simulations and our theory across several parameter values.

**Figure 5:**
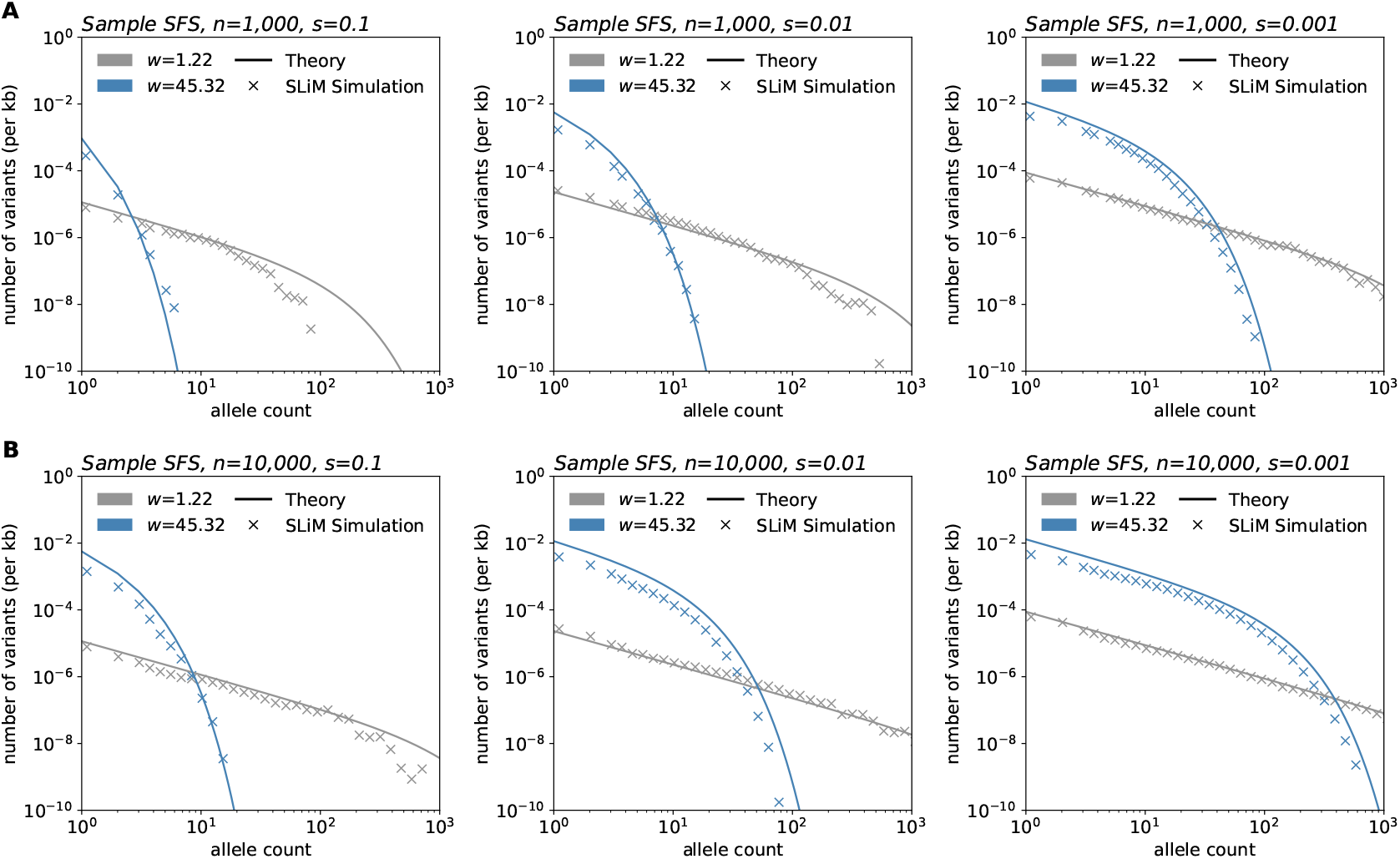
Simulations in SLiM compared to expected results under the model. SLiM simulations were performed using a modified version of the model in Battey et al. (2020). After simulation, samples of size *n* = 1, 000 (**A**) and *n* = 10, 000 (**B**) were taken at varying sampling widths. Simulations shown were run using a habitat of width 75 units, population density of 5 individuals/unit squared, and deleterious mutation rate of 10^−10^ per basepair per generation. Frequency spectra shown are averaged over 100 sampling iterations. Theory parameters are directly matched to those of the simulations.

These comparisons also reveal how computational efficiency varies greatly among the approaches. SLiM simulation time ranged between 7.58 and 11.89 hours (average: 8.64 hours) per replicate for a habitat of length 75 units (50 replicates run per sample size and selection coefficient pair). On average, the branching process simulations completed in 18.59 minutes, 1.22 hours, and 2.85 hours per one million time steps for *s*=0.1, 0.01, and 0.001, respectively for a habitat size of 10,000 units. In contrast, the time to generate theoretical frequency spectra shown in Fig. 3 ranged from 6.88 to 9.91 milliseconds per curve.

### Re-sampling experiments in the UK Biobank reveal evidence of discovery and dilution effects

Having identified relationships between the spatial breadth of sampling effort and observed variant frequencies in our theoretical work, we now consider to what extent these patterns are present in human genetic data. We artificially mimic sampling designs that vary in sampling breadth via *in silico* sub-sampling (*n* = 10, 000) individuals from the large (*n* = 469, 835) exome sequencing dataset of the UK Biobank, using sequencing data from Chromosome 1 (Backman et al., 2021). For each mock sample, we computed the frequency spectra as well as derivative summary statistics (the number of variant sites and singletons, allele frequency for variant sites and all sites, heterozygosity, and cumulative MAF).

We constructed samples spanning across two scales: fine-scale geographic sampling by birthplace within a genetically similar group (Fig. 6A-D, Fig. S16) as well as a broader sampling of genetic space as determined by the top two prinicipal components (Fig. 6E-H). On both scales, we compute distances from a center location and construct Gaussian samples with varying breadth, from highly localized near the centroid to fully uniform, adjusting for underlying heterogeneity in sampling density in UKB (Fig. S12). We label the resulting sampling designs A - H with A being narrowly centered on a location in Britain and H representing the other extreme of sampling uniformly across the full genetic space represented by the UKB cohort.

**Figure 6:**
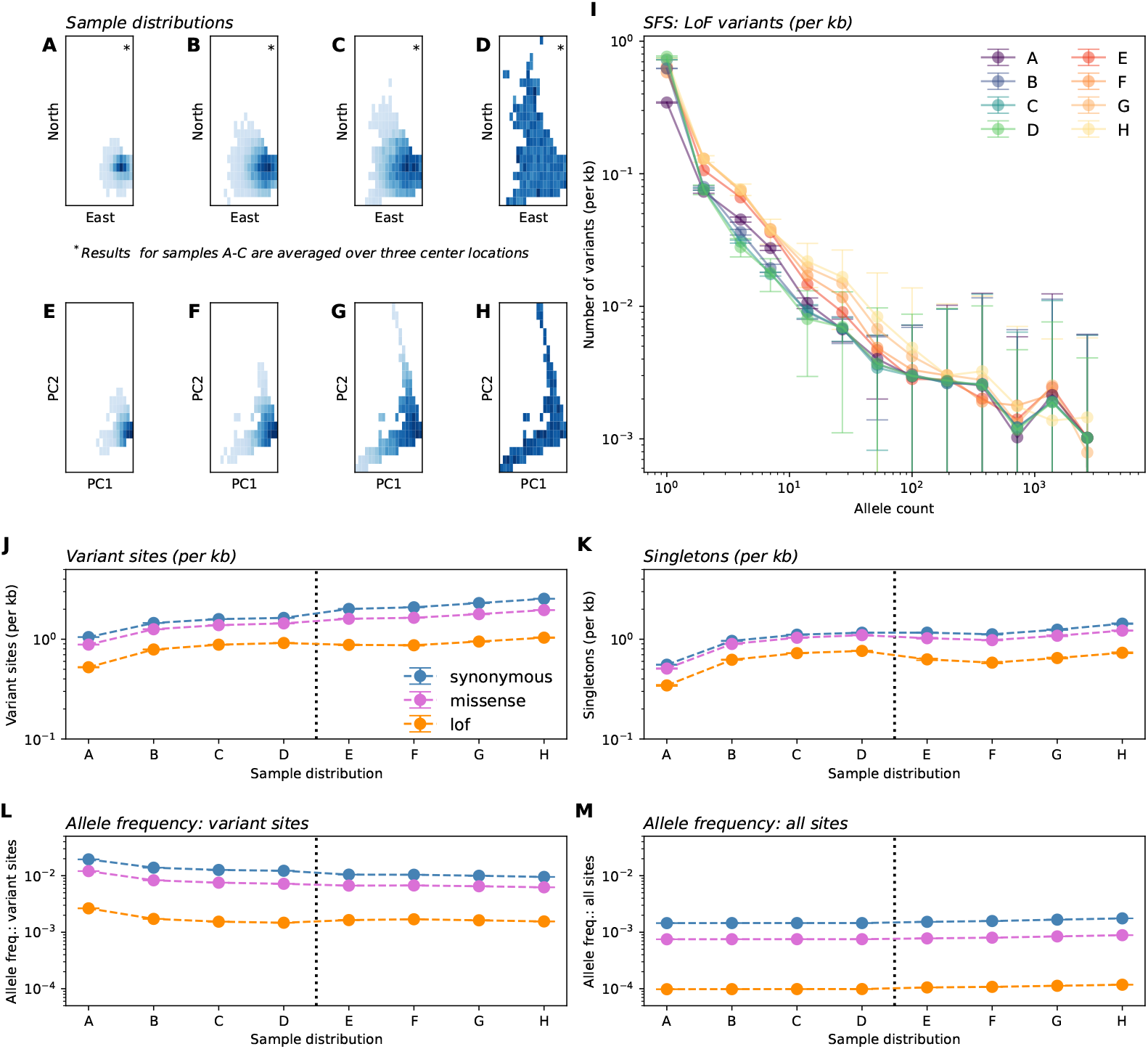
In-silico re-sampling experiments in the UK Biobank exome data. Panels **A**-**H** depict eight sample distributions. Panels **A**-**C** depict Gaussian samples in Geographic space (by birthplace) for individuals born in the UK and having similar genetic ancestry (as determined by distance in PC1-PC2 space). Sampling standard deviations, in order, are 50km, 100km, and 150km. Gaussian sampling was repeated over three center locations (see Supplemental Fig. S16) with averages across centers shown in other panels. Panel **D** depicts uniform sampling in this scheme. Similarly, panels **E**-**H** depict Gaussian samples (of the entire dataset) centered at the median in PC1-PC2 space with standard deviations 0.0015, 0,0025, and 0.005 units of Euclidean distance in PC1-PC2 space, with panel **H** depicting uniform sampling. (**I**) Average sample SFS for LoF variants on chromosome 1 across sampling distributions. (**J**-**M**) Variation in summary statistics across sampling distributions and variant annotation. All results are averaged over ten sampling replicates with error bars representing one standard deviation.

The results show patterns qualitatively similar to our theoretical predictions in that broader samples show higher proportions of variant sites overall as well as more variants on the low end of allele count (Fig. 6I-M). We find that as sampling scale becomes broader, one generally observes more variant sites, more singletons, and lower frequencies at variant sites, while mean frequency across all sites is largely insensitive to changing sampling breadth (as well as mean heterozygosity and cumulative MAF; Fig. 6J-M, Fig. S15). The differences across sampling design are larger for the putatively more deleterious loss-of-function variants, than for non-synoymous or synonymous variants.

The scale of the effects is such that for samples of size *n* = 10, 000, moving from panel A to panel B leads to on average 50.68% more discovered LoF variants with a 34.97% reduction in the sample frequency at variant LoF sites. Moving from panel A to panel H leads to 97.90% more LoF variants with a 41.36% reduction in the sample frequency at variant LoF sites. The effects of increasing sampling scale seem to moderate after reaching scale D. The mean allele frequency (across all sites) varies negligibly across the sampling scales (e.g., Panel D vs Panel E, 1.0 × 10^−4^ vs. 1.1 × 10^−4^ in LoF variants; SFS in Fig. 6I).

A deviation from theoretical predictions is that we observe a convergence of the empirical SFS across different sampling strategies for larger allele counts (e.g. allele counts greater than 10^3^; Fig. 6). We speculate this is due to the recent common ancestry of all humans, which has led to variants with large counts in any one population to be broadly shared on average (e.g., Biddanda et al., 2020), and thus such variants are plausibly less affected by sampling breadth.

## Discussion

Here, we have addressed the question of how the geographic “breadth” of sampling effort in genetic studies impacts the discovery of rare, deleterious genetic variants using a novel theoretical approach. Our analysis shows how sampling affects the expected site frequency spectrum via both discovery and dilution effects: geographically broad samples will find a greater number of variants, often at ultra-rare frequencies (e.g. singletons), and with expected counts that decay more quickly as allele frequency increases. In contrast, geographically narrow samples will include fewer variants, though these variants will appear concentrated in the sample, often at frequencies above what they would be found in uniform samples.

In several ways, our results echo the impacts of sampling on neutral variation: spatially broader samples tend to discover more variant sites overall; however, these alleles tend to be singletons and other low-frequency alleles (De and Durrett, 2007; Ptak and Przeworski, 2002; Arunyawat et al., 2007; Städler et al., 2009; Battey et al., 2020; Gloss et al., 2022). However, using our approach we can directly account for and vary the strength of negative selection, and we see that this has significant effects on predicted frequency spectra (Fig. 3, Fig. 5, Figs. S6-S9) and summary statistics (Fig. 4) for selected alleles. In particular, our analysis reveals that the more deleterious a class of variants is, the smaller the spatial scale of their spread (ℓ_*c*_) will be. In turn, we expect the effects of increasing sample breadth to saturate most quickly for more deleterious variants. That is, for more deleterious sites, the discovered alleles will be as diluted as they would be in a fully uniform sample at a comparatively smaller scale of sampling.

An unexpected result from our theory is that expected allele frequency (and in turn expected heterozygosity and cumulative MAF) is approximately constant with respect to sampling breadth. The result appears to hold in simulations and generally across several annotation categories in our empirical analysis of the UK Biobank, suggesting it is a real phenomenon in the models and data considered here.

Our results have important implications for two major areas of research that use observations of rare, deleterious variants: (i) genetic association studies and (ii) evolutionary genetic inferences of fitness effects. In the next paragraphs, we discuss our results within these two contexts.

In genetic association studies of disease phenotypes and complex traits, observed frequencies are intrinsically tied to statistical power. GWAS power is roughly linear in allele frequency for low frequency alleles, as *x*(1 − *x*) ≈ *x* for small *x*, and the cumulative MAF that impacts power in burden tests is also directly dependent on the average allele frequency. While many studies have considered the impact of increasing sample size on power, our results suggest new and interesting trade-offs related to geographic sampling breadth. Broader samples will detect a greater number of variants due to the discovery effect – and thus expand the space of potentially identifiable associations. However, each variant will have lower observed frequency (dilution) which hinders power to detect associations in single-variant GWAS designs. For instance, with a sample size of 10,000, our experiments show broad re-samples of the UK Biobank have on average 97.9% more variant sites (and 112.46% more singletons, for LoF variants), but 41.36% lower variant frequencies than when samples are narrowly concentrated (Fig. 6).

Somewhat surprisingly, these outcomes seem to largely compensate each other. In our theoretical model, the compensation is perfect, and remarkably, the average allele frequency across all deleterious sites remains constant as a function of sampling breadth. This suggests sampling scale may have negligible impact on power to detect phenotypic associations.

Such implications are tentative though – more in depth analyses of the impacts on GWAS and burden test power are needed which consider factors not addressed by our model (for instance, linkage disequilibrium patterns, corrections for population stratification, the increased rate of cryptic relatedness in narrow samples, and the effects of recent human population growth). Furthermore, the question of how best to construct samples for human genetics research is intrinsically linked to discussions of equity and inclusion in biomedical research (Bustamante et al., 2011; Popejoy and Fullerton, 2016; Dolan et al., 2023). So, while the work here contributes insights on the impact of sampling on the SFS of discovered variants, we emphasize that sampling is only one element of the multifaceted challenge of study design in human genetics.

A second area of research for which our results have key implications is the inference of fitness effects in evolutionary genetics. Many studies aim to infer the DFE from the frequencies of observed variants of different classes (Williamson et al., 2005; Boyko et al., 2008; Gutenkunst et al., 2009; Kim et al., 2017). Such studies often focus on the population-scaled selection coefficient (commonly, *Ns*) as the parameter of interest. Empirically, population genetic samples are typically taken from one or a few distinct locations, yet are modeled as a random sample from the total population. Our results imply that this practice will lead to biases in the inference of selection coefficients which will tend towards under-estimating the strength of negative selection.

Specifically, we expect that sampling narrowly from a particular location will lead to artificially high (or “concentrated”) frequencies of deleterious variants (e.g. observe how the frequency spectra becomes flatter for more narrow samples; Fig. 3A,B). In terms of our theory, this corresponds to our result that for spatially concentrated sampling, the effective selection intensity, *γ*_*E*_, can be substantially less than *Ns* (e.g. orders of magnitude lower within our test settings; Fig. 2). Thus, the frequencies used in the inference framework will be higher than expected under random mating, leading to biased estimates of *s*. This bias is likely to be most prominent for alleles under stronger selection, as the deviation of *γ*_*E*_ from *Ns* will be larger (Fig. 2). We also expect to see a downward bias in the inferred variance of the DFE for spatially concentrated samples: estimates for variants with stronger fitness effects will be biased more strongly than those with weak effects, leading to an overall reduction in variability among inferred effects.

For both of these downstream applications, another relevant finding from our model is that the magnitude of effect of sampling width on allele frequencies and summary statistics is highly dependent on the sample size (Fig. 3, Fig. 4B, Fig. S5). Thus, as sample sizes in genetics continue to grow towards millions of individuals, we may expect the impact of sampling breadth to become more evident.

The most important caveat of our work is that we analyzed a highly abstract model of a spatial population and sampling effort. While we define sampling and spatial dynamics under our model in relatively simple terms aimed to help refine one’s thinking about this problem, the realities of study designs and demography are far more complex. For instance, our model assumes that there are no boundaries on where carriers can disperse and as a result, no “boundary effects” are present. Additionally, our simple model of migration via local diffusion does not account for repeated layers of long-range dispersal events which are plausibly frequent in human and non-human populations. As a result, a geographically “narrow” sample in real data (e.g. sampling a city like London) may not truly be “narrow” in the sense of our model. We have also not considered various departures from equilibrium such as variable patterns of recent population growth, recent admixture from diverged lineages (e.g. archaic hominids), and recent origins from a shared ancestral population (e.g. shared African origins of humans). Thus, especially for settings beyond the UKB, the relevance of these more complex factors should be kept in mind.

Overall, in real studies of populations of humans or other organisms, the patterns of movement and of sampling may greatly deviate from what we investigated here. Nonetheless, the general alignment of our empirical and theoretical results suggest the real-world importance of our results for interpreting the outcomes of existing studies and designing future ones.

## Acknowledgements

We thank Jennifer Blanc, Jeremy Berg, Maryn Carlson, Castedo Ellerman, Yuval Simons, and Matthias Steinrücken for helpful comments and discussion. This work has been supported by the National Science Foundation (DGE1746045 to MCS) and the National Institutes of Health (R01 GM132383 and R35 GM149521 to JN). This research has been conducted using the UK Biobank Resource under Application Number 88057.

## 1 Supplemental Information

### 1.1 Extended theoretical methods

Here, we provide a detailed description of our theoretical methods. We model the movement, reproduction, and death of the carriers of a rare deleterious allele. These carriers are generated by mutations in a much larger population of wild-type individuals. By explicitly modeling only the rare carriers, we can approximate the evolution of their spatial distribution as a *superprocess*. We use the recent results of Friesen (2023) to find the equilibrium moment generating functional of the spatial distribution of deleterious alleles at mutation-selection-migration-drift balance.Then, we apply a model of spatial sampling to compute the expected site frequency spectra for varying spatial sampling schemes.

In the following, we will assume that all parameters (i.e., population density, mutation rate, selection coefficient, and dispersal diffusion coefficient) are spatially and temporally homogeneous. It is straightforward to specify a more general model and to apply the same procedure we outline below to calculate its site frequency spectra. However, the numerical calculations become significantly more complicated. Therefore, we focus on getting intuition from the scaling results in the homogeneous case and leave generalizations to future work.

#### 1.1.1 Population genetic model

We consider a population of organisms living in a habitat *H*. Here we will focus on 2 − dimensional continuous habitats, so that *H* ⊂ ℝ^2^, but most of the theory applies to more general habitats. For simplicity, in our numerical calculations we will use a toroidal habitat of length *L*, i.e., *H* = [0, *L*]^2^ with periodic boundary conditions. Let *ρ*_*N*_ be the population density measure so that the number of individuals living in a region *A* ⊂ *H* is given by 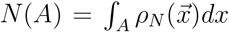. The total population size is 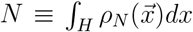. We assume that *ρ*_*N*_ is large and stably maintained by ecological forces so that we can neglect random fluctuations in population density due to migration, births, and deaths.

We are interested in tracking the number and spatial distribution of carriers of a rare deleterious variant with fitness cost *s* > 0. We will restrict ourselves to the weak-selection regime, *s* ≪ 1. With this assumption, carriers of the deleterious allele have 1 − *s* offspring on average, compared to 1 for non-carriers. By focusing on rare alleles, we can neglect dominance effects because homozygous carriers should make up a negligible fraction of the carriers. We assume that wild-type alleles undergo mutation to the deleterious allele at rate *µ* per generation per individual. Thus, we model the influx of *de novo* mutations as a Poisson point process with intensity 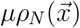. This approximation neglects the reduction in the mutation supply due to the fact that some fraction of the *ρ*_*N*_ might already carry the mutation. However, if *µ* ≪ *s*, the effect of selection dominates the resulting reduction in mutation supply and we can neglect it. We will similarly neglect back-mutation from the mutant to the wild-type.

For mathematical tractability, we will assume that carriers of the deleterious allele reproduce, die, and move about the habitat independent of one another and of the background of wild-type individuals. This assumption is justified as long as the deleterious allele remains rare. Given the rare-allele assumption, we model the movement of an individual carrier as a continuous-time Markov process on *H* with infinitessimal generator *σ*^2^∇^2^ (i.e., translation invariant, isotropic diffusion), independent of the positions of other carriers. This simplification is similar to Haldane’s branching process approximation to the Wright-Fisher process (Haldane, 1927), and has since been applied in spatial population genetic modeling (for instance, Novembre and Slatkin, 2009).

In a continuous habitat we need to choose a suitable definition of ‘rare’ that accounts for spatial variation in the mutant frequency. To wit, let *σ* be the average distance an individual will move in one generation (more formally, the root mean squared difference between parent and offspring birth locations). We assume that the number of carriers in a ball 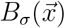 with radius *σ* about any point 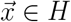 is small compared to 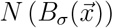 , Wright’s neighborhood size (Wright, 1946). As *σ* is the shortest length scale in the model, it is reasonable to assume that fluctuations in allele frequency at distances shorter than this scale will be short-lived and not contribute to the long-term evolution of the population.

Having defined the mutational process by which carriers are generated and the migration process by which they move around the habitat, it remains to define the mechanism by which they die and reproduce. Consistent with our assumption that carriers behave independently, we will model reproduction as a continuous-state branching process. Combining this with our Markov process for movement yields a superprocess model of the evolution of the spatial distribution of carriers (Watanabe, 1968; Dawson, 1993). For overviews of superprocesses and their properties, see Le Gall (1999); Etheridge (2000). If we add an influx of particles according to the mutational point process with intensity *µρ*_*N*_ , we have a superprocess with immigration (Kawazu and Watanabe, 1971). (Confusingly, in the superprocess literature, our mutation process is called “immigration” and the migration process is sometimes called “mutation”.) We now define this process and then give the main result (Eqs. 10–12) needed to calculate sample properties.

A superprocess {*Z*_*t*_}is a measure-valued random process. That is, {*Z*_*t*_} is a set of measures on the habitat *H* indexed by time, *t*. Measures are defined by how they integrate functions over their domain. Accordingly, we introduce the inner product ⟨*Z, f* ⟩ ∈ ℝ, defined as:

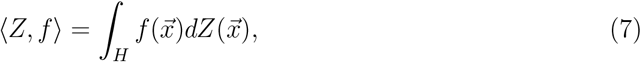

where *Z* is a finite measure on *H*, and *f* : *H* → ℝ is a measurable function on *H* (Le Gall, 1999). The probability distribution of a superprocess at time *t* is characterized by its *moment generating functional* (MGF), Φ_*t*_, defined as:

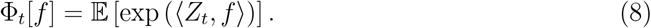

Just as the derivatives of the moment generating function of a random variable gives its moments, the functional derivatives of Φ_*t*_ with respect to *f* give moments of the inner product ⟨*Z*_*t*_, *f*⟩.

In our model, *Z*_*t*_ measures the number of carriers in a region of space. For a region *A* ⊆ *H*, we define an indicator function 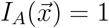 for 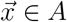 and zero otherwise, so that:

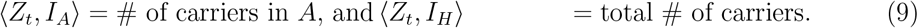

By analogy, ⟨*Z*_*t*_, *f*⟩, for an arbitrary non-negative *f*, gives the counts of carriers according to their positions by the weighting function *f*.

Friesen (2023) has recently shown that subcritical superprocesses (i.e., ones where the measure decays exponentially) with immigration tend to a stationary distribution, subject to various technical conditions. In particular, for our process with mutational supply intensity *µρ*_*N*_, diffusion coefficient *σ*^2^, and selection coefficient *s*,

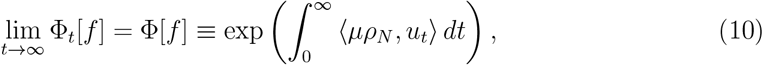

where *u*_*t*_ is the solution to the semilinear PDE:

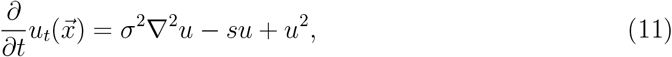

subject to initial condition:

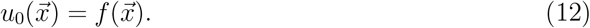

The function *u* does not have a direct biological interpretation, though its associated PDE incorporates several features of the evolutionary process (including selection and drift).

The stationary MGF, Φ, completely characterizes the counts and spatial distribution of carriers of the deleterious alleles of a population at steady-state. These patterns are due to the balance between the forces of mutation, selection, genetic drift, and migration. In the rest of this section, we will show how Eq. 10 can be used to calculate the expected site frequency spectrum for a spatially localized sample from the population.

#### 1.1.2 Spatial sampling the allele frequency distribution

We now connect the superprocess model of allele frequencies to the site frequency spectrum of a finite sample. We are interested in geographically biased samples, where the probability that an individual is sampled depends on its location according to a *sampling density*. Let the kernel function *g*(·) be the shape of the sampling density so that *∫*_*H*_ *g*(*x*)*dx* = 1. To capture the effect of broad versus narrow sampling, we will consider sampling kernels with a scale parameter *w*, which represents the typical distance between sampled individuals. In particular, for sampling on a torus, we will use a bivariate wrapped Gaussian sampling kernel with standard deviation *w*.

Note that the sampling density is a measure of the *a priori* sampling effort across space, rather than the realized locations of sampled individuals. In the following, we assume that we do not have access to the locations of our sampled individuals. However, this framework could be extended to consider the joint statistics of samples taken from multiple known locations.

We define the expected value of the *k*’th element of the site frequency spectrum (SFS) as the expected fraction of sites with *k* copies of the deleterious allele 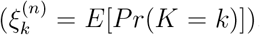. Assuming that the samples are taken independently from the population with replacement, the number of copies of the deleterious allele is the sum of n Bernoulli trials with a random probability of success:

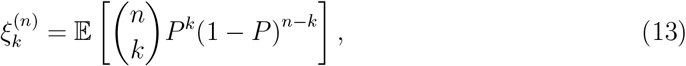

where *P* is a random variable representing the probability any particular sampled allele is deleterious.

The distribution of *P* depends on (1) the locations of carrier individuals characterized by the superprocess *Z*, and (2) the probability that we sample a carrier given its location. For a sample taken uniformly from a region *A* ⊆ *H, P* is the fraction of carriers in *A*:

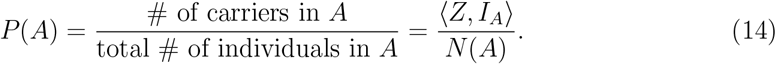

If, instead, the sample is taken by choosing among a countable set of regions {*A*_*i*_} according to probabilities {*g*_*i*_} and then sampling uniformly within the chosen region,

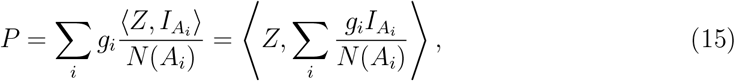

where the second equality uses the linearity of inner products. Taking the limit that sampling probabilities vary continuously according to a localized sampling density, we have

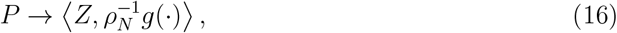

so that the sampling probability is an inner product of the random measure *Z* with the population-scaled sampling density.

Therefore, using Eqs. 8, 10, and 16, the moment generating function of *P* at steady-state is given by

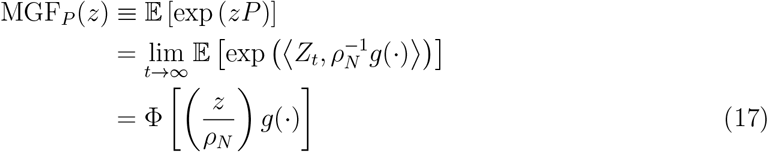

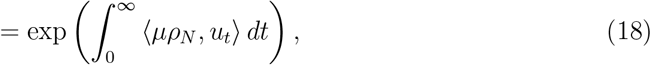

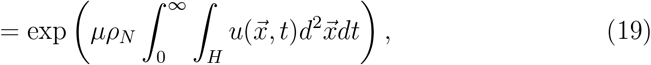

where *u*_*t*_ solves Eq. 11 with initial condition

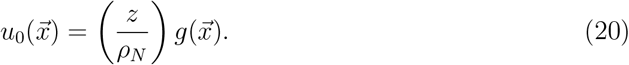

In the following sections, we will use these equations to solve for key properties of the distribution of *P* in order to evaluate the SFS per Eq. 13.

#### 1.1.3 Moments of the allele frequency distribution

Here, we aim to calculate the log of the MGF of *P* by way of Eq. 19. We consider a habitat *H* = [−*L*/2, *L*/2]^2^ (for *L* ∈ ℝ) with periodic boundary conditions. For mathematical convenience, we will work in Fourier space. First, we take a Fourier transform of 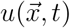 over both time and space:

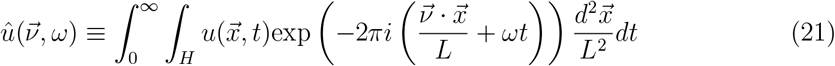

where *ω* ∈ ℝ (temporal frequency) and 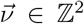(spatial frequency). Applying the same transformation to the PDE in Eq. 11 gives:

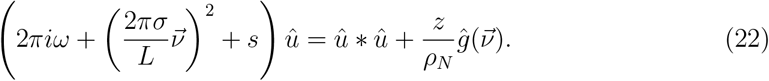

It then follows from Eqs. 19 and 21 that:

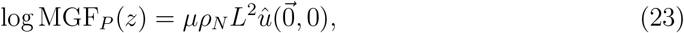

and as such we aim to solve for the value of *û* at the origin.

To proceed, we calculate a perturbative expansion of *û* in powers of *z*, up to the second order:

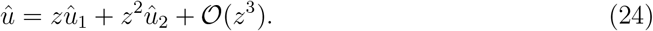

Substituting into Eq. 22 gives:

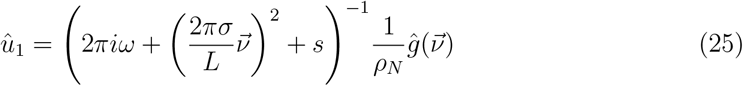

and

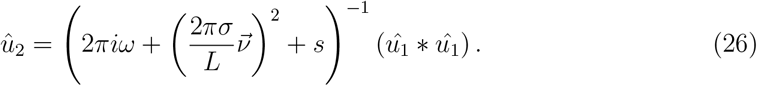

Evaluating at the origin, we have:

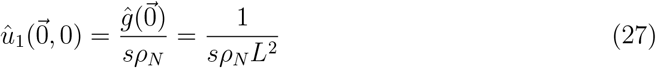

and

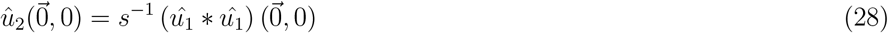

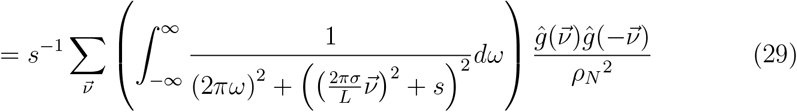

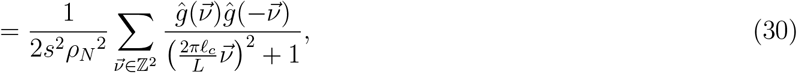

where the last line introduces the critical distance 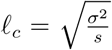. We refer to this value as the *characteristic length scale* (see section 3.1 in the main text).

From here, we can calculate the mean and variance of *P*. We find that the mean allele frequency is independent of the sampling scheme:

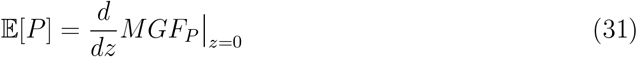

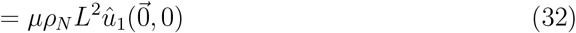

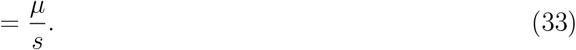

On the other hand, the variance is given by:

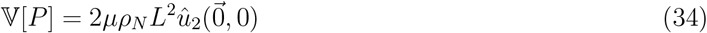

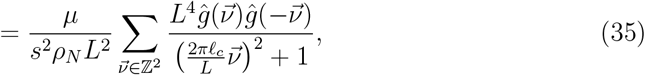

and so is dependent on the sampling kernel *g*(·). If sampling is uniform over the habitat,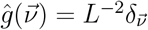. Thus,

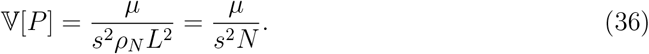

On the other hand, if sampling follows a wrapped normal distribution with scale *w*,

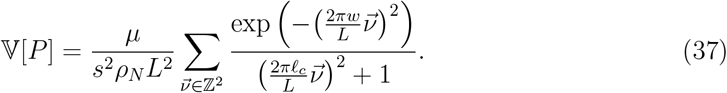

For *w* ≳ *L*, the numerator of the sum falls off rapidly with 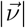 and we converge to the uniform sampling result. For 𝓁_*c*_ ≫ *L*, the denominator grows large for 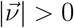 and we again converge to the uniform sampling result. For *w*, 𝓁_*c*_ ≪ L, we can approximate the sum with an integral:

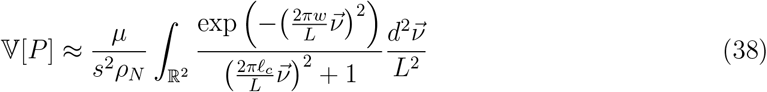

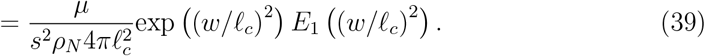

For *w* ≫ *l*_*c*_, this is approximately *µ*/(*s*^2^*ρ*_*N*_ 4*πw*^2^), which implies that we converge to the uniform sampling result when 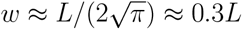 For *w* → 0, the expression diverges as we integrate over very large frequencies, which correspond to very small length scales where our model breaks down. We can remedy this by imposing a cutoff on 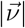. The smallest length scale our model can sensibly talk about is *σ*, the per-generation dispersal distance. Thus, a reasonable cutoff is 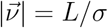.

We validate these expressions by comparing results for 𝔼 [*P*] and 𝔼 [*P*^2^] in simulations to their respective values under the model, and find a close correspondence (Fig. S4). Additionally, as the sampling width approaches the habitat width in the simulations, values of 𝔼 [*P*^2^] approach expected values under uniform sampling, as expected.

#### 1.1.4 Effective parameters and the sample SFS

Having calculated the first two moments of *P*, we will now use them to approximate the full distribution of *P* and use this to calculate the SFS of a sample of finite size. We will assume that *P* approximately follows a Gamma distribution whose parameters we can calculate from the first two moments. If sampling is uniform over the habitat, it can be shown that this holds exactly (see section 1.1.6), and we show via simulation that this assumption is reasonable for non-uniform sampling as well (Figs. S2 and S3). Thus, we assume *P* follows:

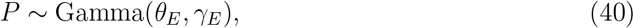

and we will refer to the shape and rate parameters, *θ*_*E*_ and *γ*_*E*_, as the *effective mutation supply* and *effective selection intensity*, respectively. The motivation behind these names will be clarified by their derivation.

We derive the form of these parameters for both the uniform sampling and wrapped normal sampling cases using the method of moments applied to previous results (Eqs. 33- 39). When sampling is spatially uniform, we have:

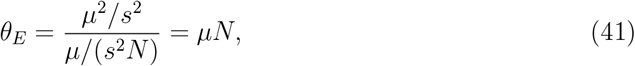

and

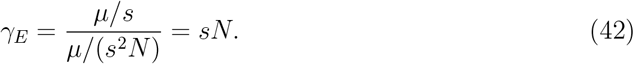

When sampling follows a wrapped normal distribution with scale *w*, we have:

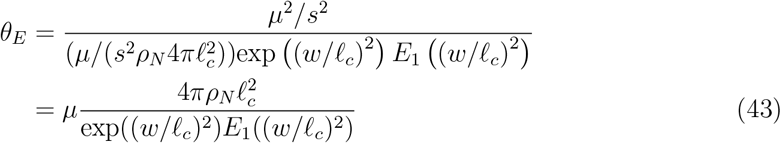

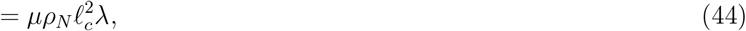

where:

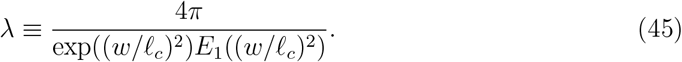

Similarly, we have:

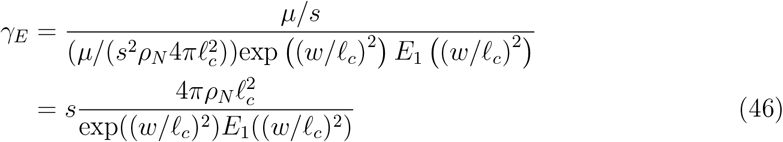

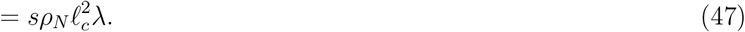

The compound scale factor λ (which we refer to as the *sampling effect scalar* ) captures all spatial sampling aspects of the problem.

To summarize, we define the distribution of *P* for the uniform sampling case as:

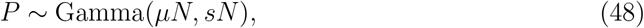

and for the wrapped Normal case as:

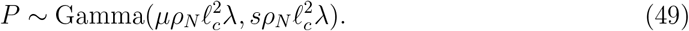

For a sample of *n* haploid genomes, let *K* ∈ {0, … , *n*} be a random variable representing the number of sampled copies of the deleterious allele in the focal site. Recall from Eq. 13 that the number of copies of the deleterious allele is the sum of *n* Bernoulli trials with probability of success *P*, or equivalently, *K* ∼ Binom(n, *P* ). For large *n* and small P, this is approximately *K* ∼ Pois(nP). Then, from properties of Gamma-Poisson mixtures, allele counts in a finite sample of size n follow:

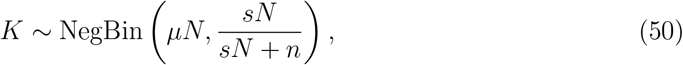

for the uniform sampling case, and:

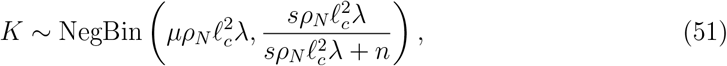

for the wrapped Normal sampling case. We can use these distribution to calculate elements of the SFS as:

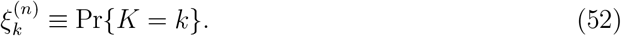

The remainder of the results follow from these expressions.

#### 1.1.5 Derivation of summary statistics

Having derived the form of the SFS, we can now obtain expressions for various population genetic summary statistics. We will show explicit derivations only in the case of wrapped Normal sampling, for brevity, though similar derivations can be obtained easily for the uniform sampling case. First, we consider the expected proportion of variant sites in a sample, or equivalently the probability that a particular allele segregates in a sample of size n. This follows from Eq. 51:

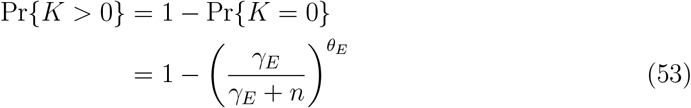

Mean allele frequency is invariant to the scale of sampling:

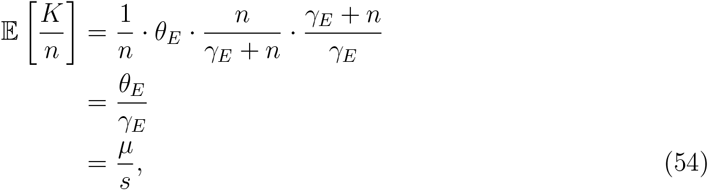

though the mean frequency of *non-monomorphic* alleles does vary according to the sampling design. We obtain an expression for the conditional mean as follows:

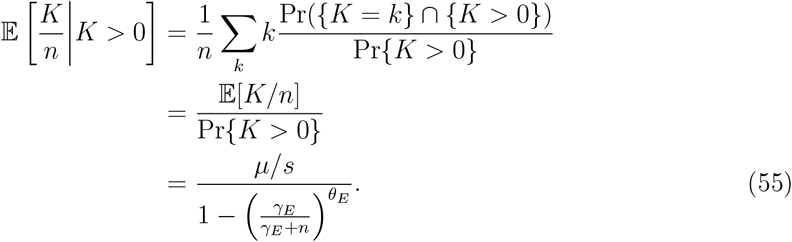

Heterozygosity is defined as the probability that two alleles are different from one another, and so it follows from the distribution of *P* rather than the SFS. Accordingly, we calculate expected heterozygosity as follows, using Eq. 40:

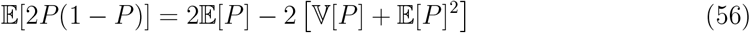

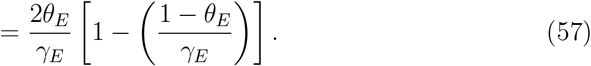

We calculate cumulative MAF, which is informative of burden test power, following the definition of Wang et al. (2014). In the context of our theoretical results, this has the form:

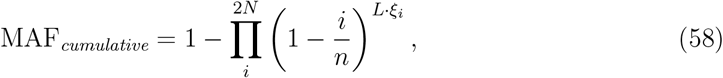

where *ξ*_*i*_ is the expected number of variants at count *i* per basepair and *L* is the length of the genomic region (bp).

We note that this expression for cumulative MAF is approximately proportional to the expected per-site allele frequency and heterozygosity. To see this, we first express the cumulative MAF in terms of 𝔼[P], for small *i*/*n* and small *L*𝔼[*P*]:

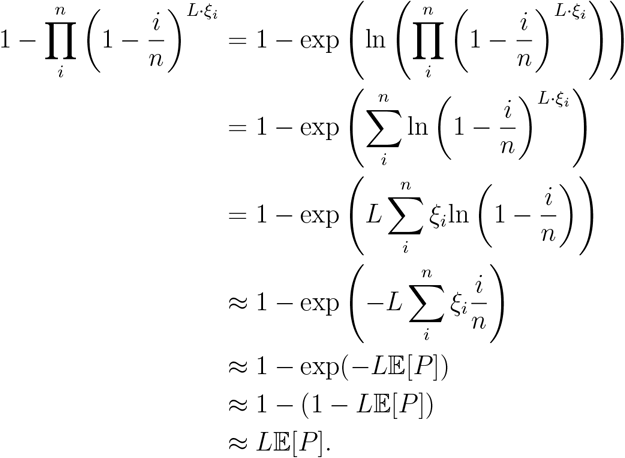

For small 𝔼[*P*], we have an expected heterozygosity of approximately 2𝔼[*P*]. This implies that the cumulative MAF is approximately proportional to expected heterozygosity, with cumulative MAF being larger by a factor of *L*/2.

#### 1.1.6 Exact solution for the uniform sampling case

Here, we provide an exact derivation of the distribution of *P* when sampling is uniform, providing motivation for our approximation that *P* is approximately Gamma-distributed more generally. Eq. 11 is a nonlinear parabolic PDE and can not be solved in closed form for general initial conditions. However, if we sample uniformly over the habitat, *u*_*0*_(*x*) becomes constant and ∇^2^*u* = 0, yielding a Bernoulli ODE for *u*, which we can solve exactly. Note that in our model, uniform sampling is equivalent to sampling from a panmictic population. This is because we are focused on rare alleles, which are assumed not to interact. For common alleles, local fixation changes the dynamics qualitatively and breaks this equivalence.

For uniform sampling, *g*(*x*) = 1/*L*^2^, and the spatial derivative term vanishes so that Eqns. 11, 19, and 20 become:

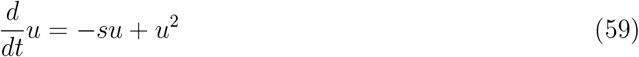

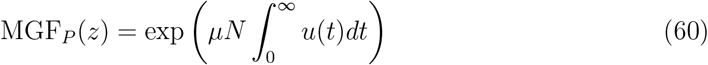

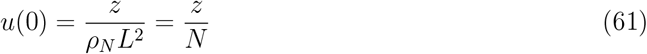

At this point, we could solve Eq. 59 directly. Instead, we will motivate our approach to the non-uniform sampling case by finding a power series solution for 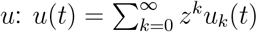. Substituting into Eq. 59 and organizing the terms by powers of *z*, we can generate an infinite sequence of ODEs for the terms {*u*_*k*_}. Starting with *k* = 0, we have

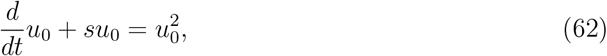

with initial condition *u*_0_(0) = 0. This has the trivial solution *u*_0_ = 0.

For *k* > 0, we have

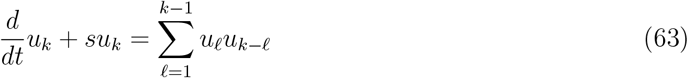

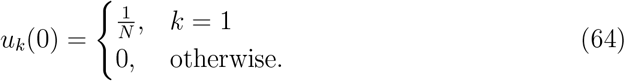

Each equation in the hierarchy is a first-order linear ODE with a forcing term that depends only on the solutions to lower-order terms.

Then it can be shown by induction that for all *k* ≥ 1, the following holds:

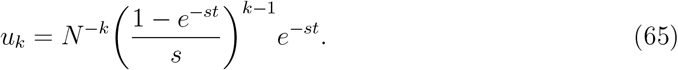

Then, by property of a geometric series:

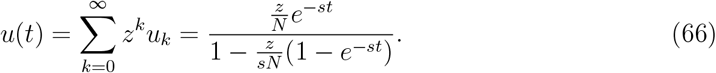

Substituting into equation Eq. 61 gives:

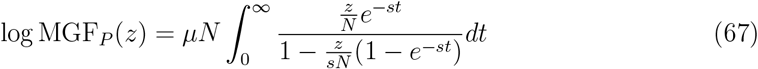

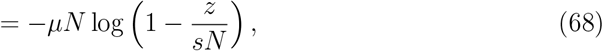

and so:

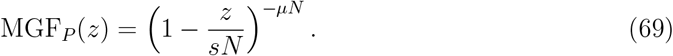

Thus, *P* follows a Gamma distribution with rate *sN* and shape *µN*, which is consistent with predictions from classical population genetic models (Wright, 1940; Kimura and Ohta, 1978). Moreover, this foreshadows our results that the form of the SFS is determined by two compound parameters representing selection and mutation, respectively.

### 1.2 Extended simulation methods

Here, we provide details on our simulation methods. All simulation code and associated scripts are available at: https://github.com/NovembreLab/spatial rare alleles.

#### 1.2.1 Spatial branching process simulations

Our first set of simulations is based on a branching process framework and aligns closely with our theoretical model. The habitat is a square of length *L* with periodic boundary conditions. Consistent with our theoretical model, carriers appear *de novo* with rate *µ* · *ρ*_*N*_, give birth with rate 1 − s, and die at rate 1. Between events, dispersal of individuals occurs according to a Gaussian distribution with variance *σ*^2^*t* where *t* is the time between events. We sample alleles at random times at rate *r*, according to a wrapped Gaussian sampling kernel with scale parameter *w*. We implement these simulations via the Gillespie algorithm (Gillespie, 1977). For computational efficiency, we implement a form of pseudo-replication in that for each simulation, we sample 100 evenly-spaced sampling centers within the habitat. Each simulation runs for 10 million generations.

Initially, we use these simulations to confirm that a negative binomial PMF provides a good fit to the simulated SFS via the method of moments (Fig. S2-S3). The ouput of each simulation is a vector of sampled values of *P*, which we use to calculate the first two moments of the allele frequency distribution: 𝔼 [*P*] and 𝔼[*P*^2^] (Fig. S4). Having computed these moments, we can then calculate the key parameters of the expected SFS under our model (equivalently to Eq. 44 and Eq. 47:

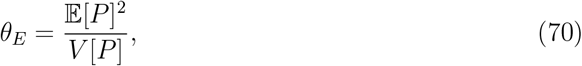

and

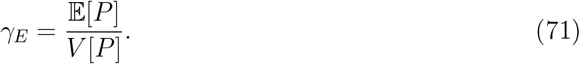

We can then use ratios between these terms to compute the λ parameter.

#### 1.2.2 SLiM simulations

Our second set of simulations implements a previously developed spatial model (Battey et al., 2020) in SLiM (Haller and Messer, 2019). All conditions are the same as in Battey et al. (2020) except that all variants are deleterious with some selection coefficient. For each selection coefficient and sample size of interest, we run 50 replicates of the simulation in a square habitat 75 units wide with a population density of 5 for a genome of length 100 Mbp and mutation rate 1 × 10^−10^ per base pair per generation. For each simulation run, we sample individuals according Gaussian (with varying width) or uniform distributions and obtain the sample SFS. We then average over 100 sampling iterations for each width.

The model from Battey et al. (2020) includes several factors that are not modeled in our theory or branching process simulations. For instance, their model includes non-toroidal boundary conditions, with the probability of individual survival declining near range edges to avoid upward biases in fitness. A particularly notable difference in the two models is the definition of the parent-offspring distribution. In our model, individuals disperse away from their location of origin (which is a single point) at root mean squared distance *σ* per generation. Under the Battey et al. (2020) model, individuals arise as the offspring of two parents, and dispersal occurs from each parent 50% of the time. This results in a constant scalar difference in all spatial parameters between the Battey et al. (2020) model and ours (under our simulation parameters, this scalar works out to 4.08). However, we find that since both w and *σ* parameters in our model are scaled by this factor, it cancels out of *w*/𝓁_*c*_ and thus λ as well as 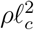. As a result, estimates of *θ*_*E*_ and *γ*_*E*_ (and thus, all downstream results) are not changed affected by this difference. We note that time-scales for mutation are the same between models (per-generation).

### 1.3 Extended empirical methods

Here, we provide additional detail on our empirical methods. Scripts are available at: https://github.com/NovembreLab/spatial rare alleles. Additional source code is available upon request.

#### 1.3.1 Sampling importance resampling algorithm and implementation

A key step in our empirical analysis is to construct samples within the UK Biobank having Gaussian or uniform distribution. Here, we provide detail on the sampling procedure used.

For samples of individuals in geographic (birthplace) space, we first filter the data to individuals passing QC (per UK Biobank metrics Bycroft et al., 2018), born in the UK, with coordinates available, and having Euclidean distance within 0.0001 of the median centroid in PC1-PC2 space. We then calculate binned frequencies of birthplaces in discretized geographic space (20×20 grid).

For uniform samples, we assign weights that are inversely proportional to the frequency of individuals in the bin in which an individual lies. For Gaussian samples, we compute distance per-individual from one of three pre-selected center points (located centrally within a bin) and assign weights according to a Gaussian density with standard deviation w, divided by the binned frequency as used in the uniform weight calculation. Centers were chosen to avoid known urban or otherwise high-density areas (to avoid model mis-specification) and such that center locations had sufficient individuals in the nearby region as to avoid extreme re-sampling of individuals. For samples in PCA space, this process is identical except that we use all individuals passing QC, construct the 20×20 grid over PC1-PC2 space, and only use one center point (which corresponds to the bin including the median value). All weights are normalized to sum to one.

Then, using custom scripts, for each set of weights we sample sets 10, 000 individuals *with replacement*. As a result of the weighting scheme used and this sampling step, the birthplace locations/PC1-PC2 coordinates for each sample match either a uniform distribution or a Gaussian distribution with weight w, as intended (see, for instance, Fig. S16). We then compute and output the SFS for each sample.

#### 1.3.2 Calculation of summary statistics

Having outputted the SFS, we then use it to compute various summary statistics. The number of variant sites, number of singletons, average variant frequency, and average heterozygosity were all calculated using standard formulas. We note that cumulative MAF was calculated following the definition of Wang et al. (2014) using the below formula:

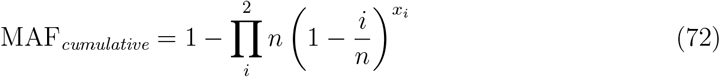

where *x*_*i*_ is the number of alleles at count i in the empirical SFS.

All results shown are averaged over ten sampling replicates each, and values are reported per-kb as appropriate. Additionally, metrics for samples within geographic (birthplace) space are averaged over each of the three sampling centers.

## 2 Supplemental Figures

**Figure S1:**
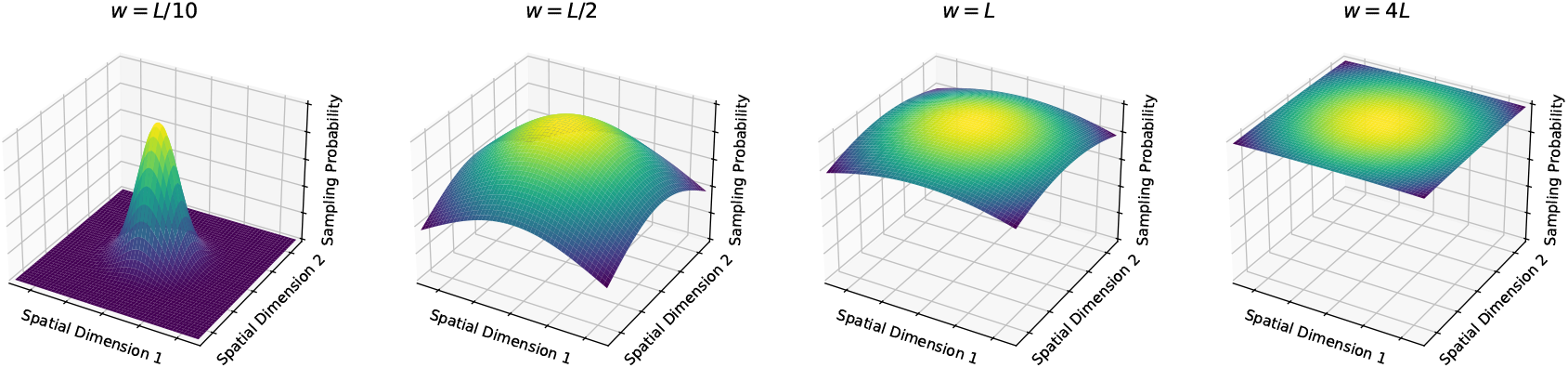
Visualization of a Gaussian sampling kernel on a square habitat of size *L* × *L*. Values of w are, ranging from “narrow” to “broad”.

**Figure S2:**
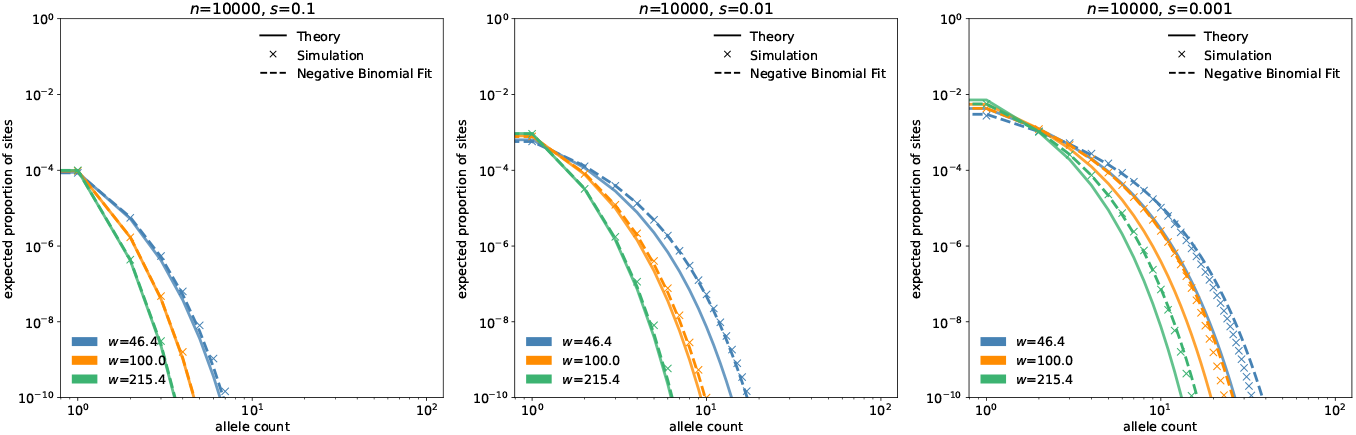
Simulated frequency spectra (x’s) with sample size *n* = 10, 000 for a range of selection coefficients. Dashed lines indicate a negative binomial PMF fit to simulation results via the method of moments. Solid lines indicate theoretical expectation. Other model and simulation parameters include: *σ* = 10, *ρ* = 20, *L*=1,000, and *µ* = 1*e* − 9.

**Figure S3:**
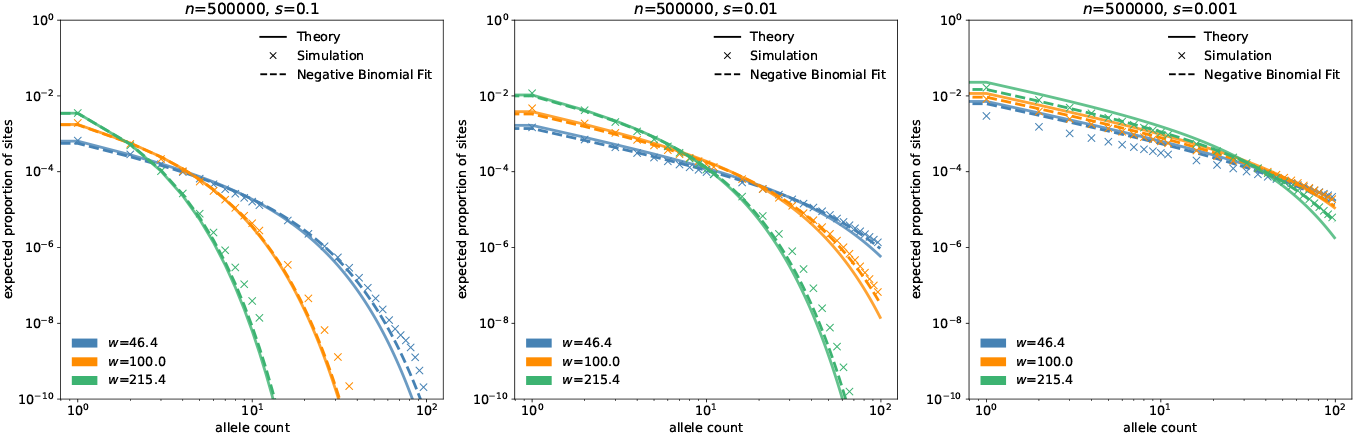
Simulated frequency spectra (x’s) with sample size *n* = 500, 000 for a range of selection coefficients. Dashed lines indicate a negative binomial PMF fit to simulation results via the method of moments. Solid lines indicate theoretical expectation. Other model and simulation parameters include: *σ* = 10, *ρ* = 20, *L*=1,000, and *µ* = 1*e* − 9.

**Figure S4:**
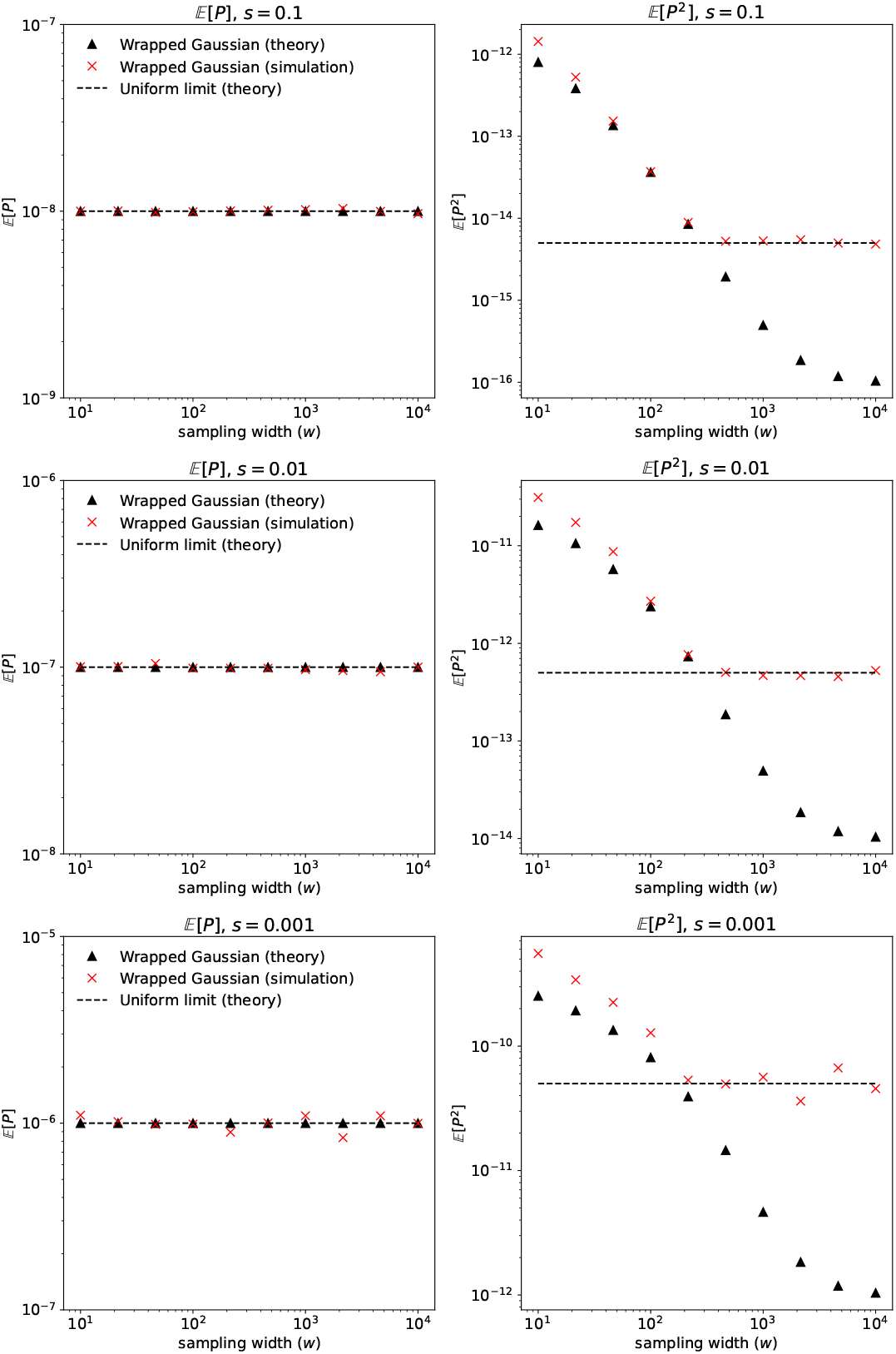
Theory and simulation results for the first and second moments of *P* over a range of sampling widths and selection coefficients. Dashed line shows expectation under uniform sampling, while other markers indicate wrapped Gaussian sampling. Other model and simulation parameters include: *σ* = 10, *ρ* = 20, *L*=1,000, and *µ* = 1*e* − 9.

**Figure S5:**
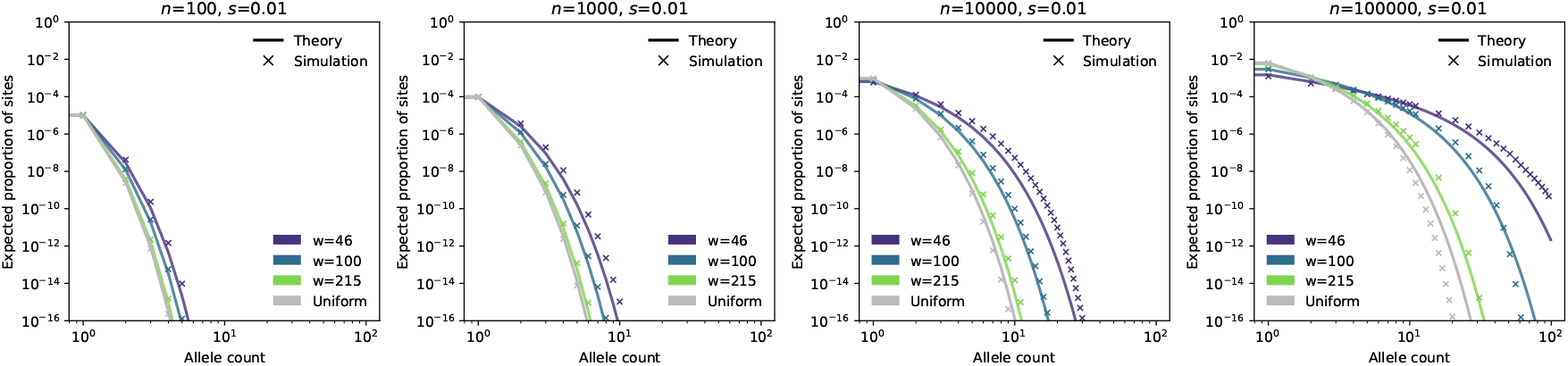
Expected site frequency spectrum from theoretical model and branching process simulations for increasing sample size (left to right). Other model and simulation parameters include: *σ* = 10, *ρ* = 20, *L*=1,000, and *µ* = 1*e* − 9.

**Figure S6:**
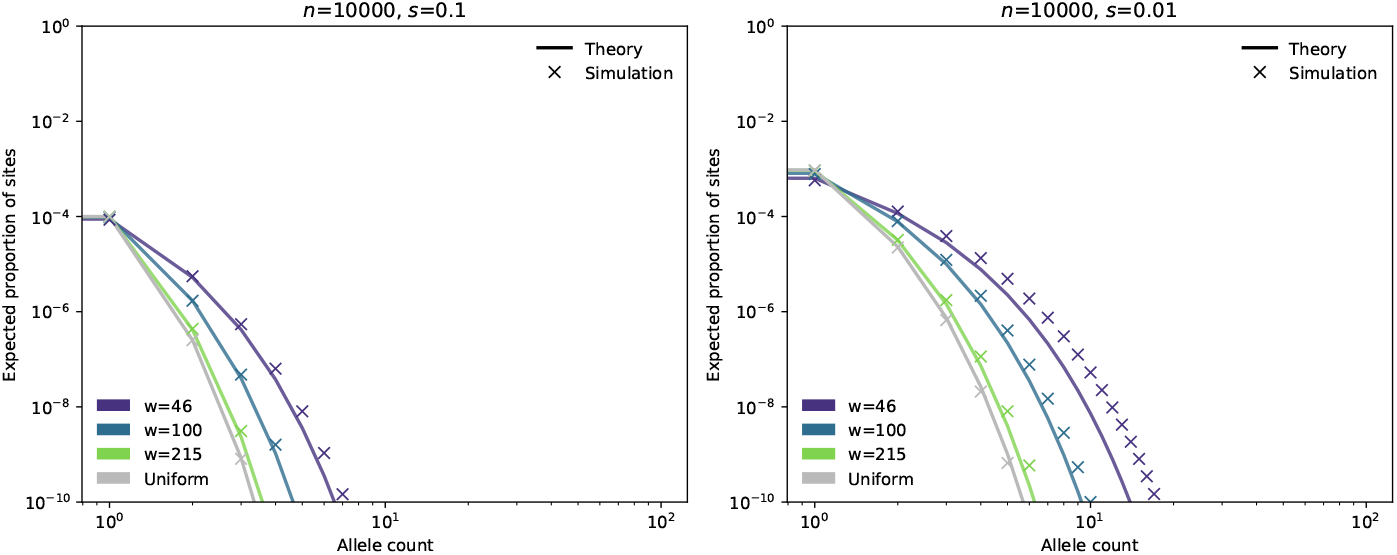
Expected site frequency spectrum from theoretical model and branching process simulations for stronger (left) and weaker (right) selection. Other model and simulation parameters include: *σ* = 10, *ρ* = 20, *L*=1,000, and *µ* = 1*e* − 9.

**Figure S7:**
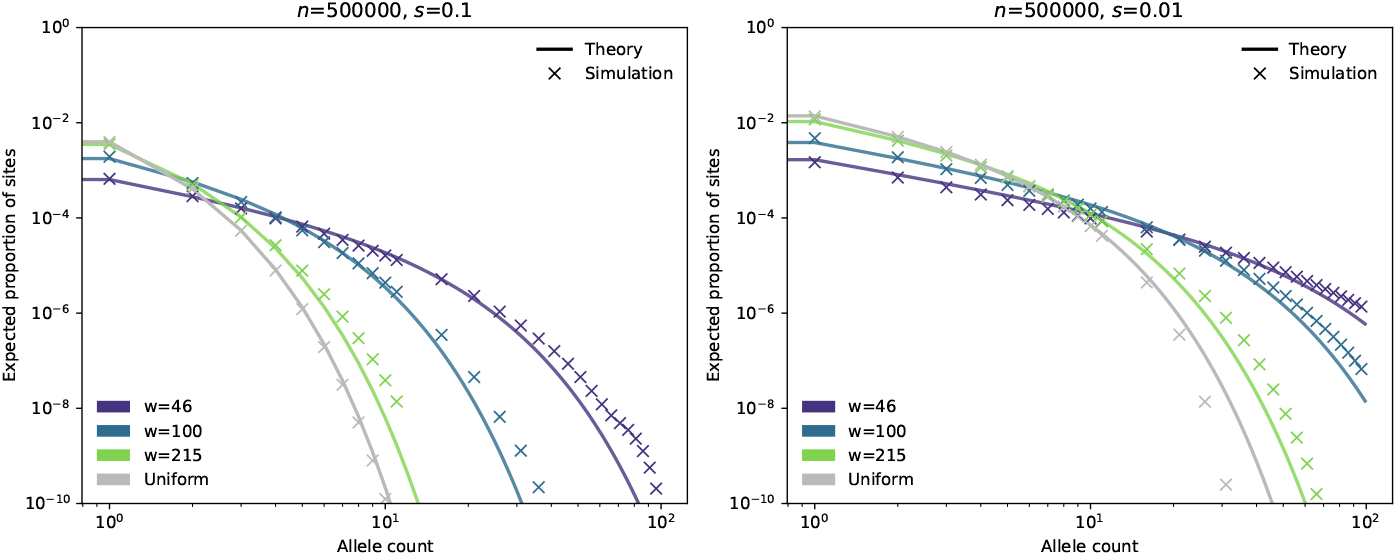
Expected site frequency spectrum from theoretical model and branching process simulations for stronger (left) and weaker (right) selection. Other model and simulation parameters include: *σ* = 10, *ρ* = 20, *L*=1,000, and *µ* = 1*e* − 9.

**Figure S8:**
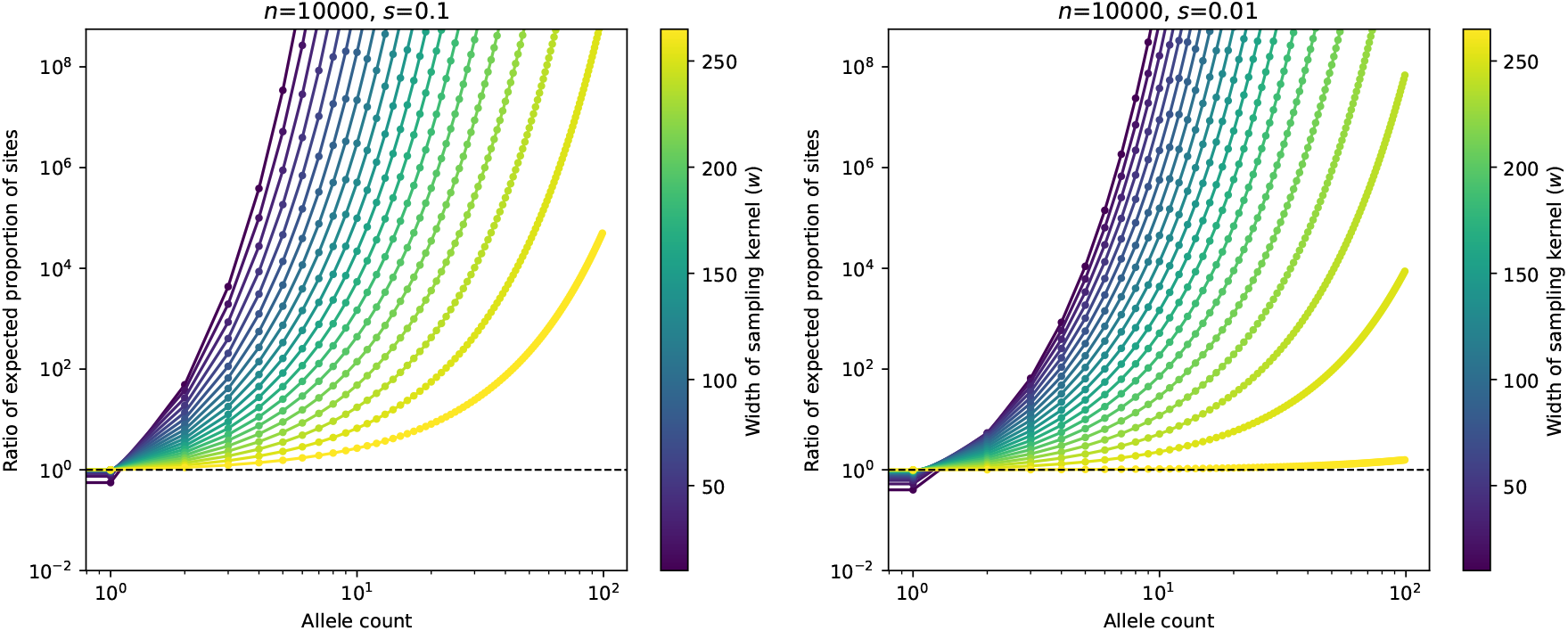
Ratio between theoretical SFS values in a sample of width *w* vs. uniform sampling for stronger (left) and weaker (right) selection. Other model and simulation parameters include: *σ* = 10, *ρ* = 20, *L*=1,000, and *µ* = 1*e* − 9.

**Figure S9:**
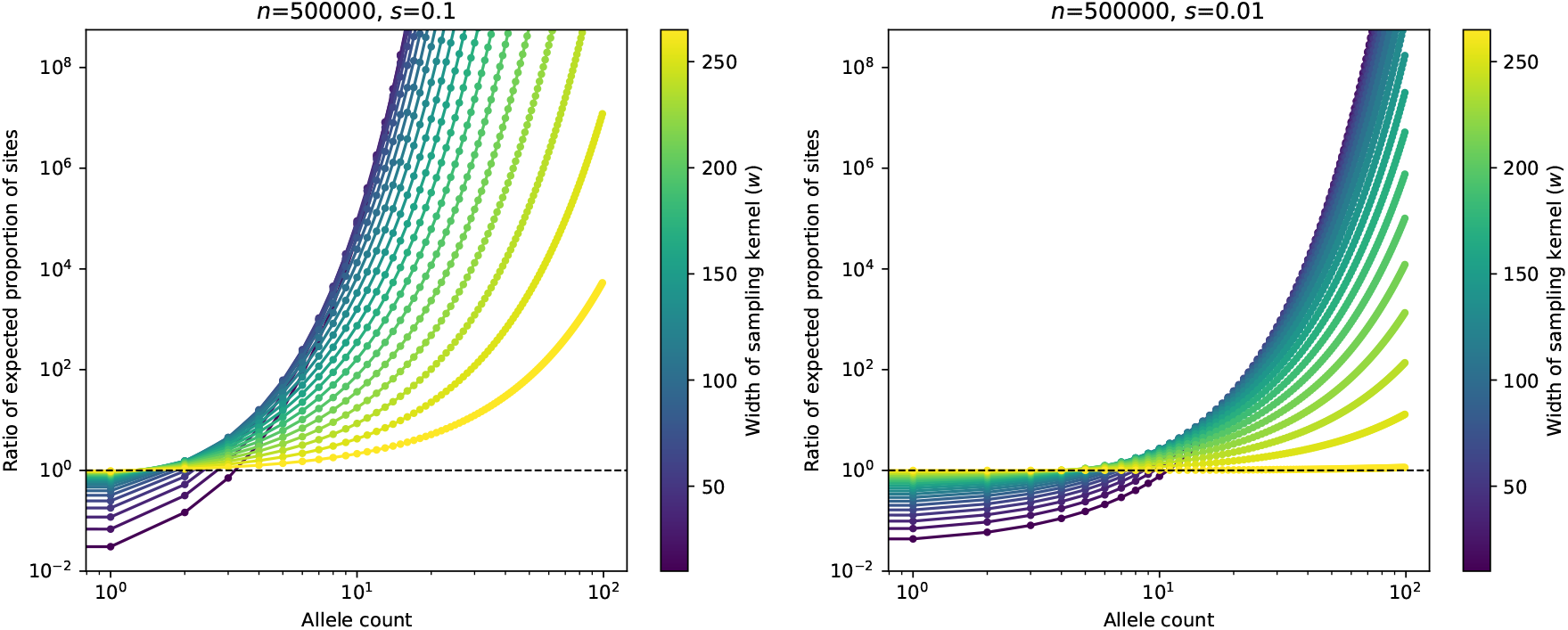
Ratio between theoretical SFS values in a sample of width w vs. uniform sampling for stronger (left) and weaker (right) selection. Other model and simulation parameters include: *σ* = 10, *ρ* = 20, *L*=1,000, and *µ* = 1*e* − 9.

**Figure S10:**
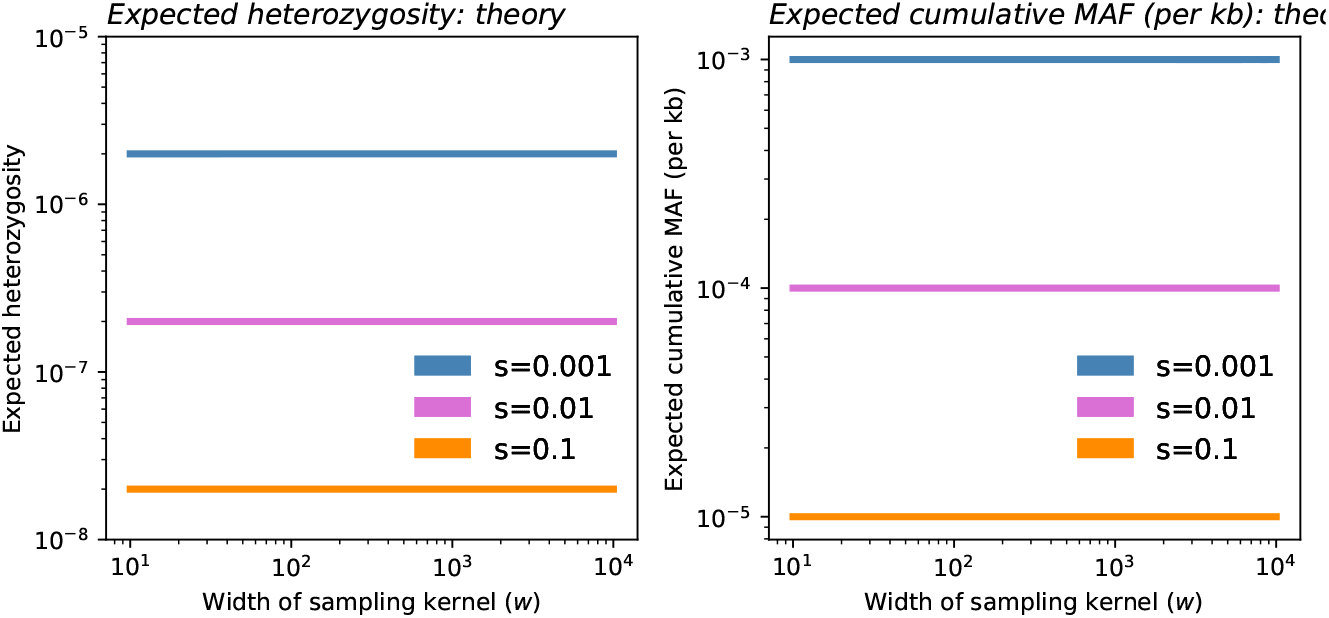
Expected heterozygosity and cumulative MAF (per kb) from theory. Inplots shown, *σ* = 10, *ρ*_*N*_ = 20, and *µ* = 1*e* − 9.

**Figure S11:**
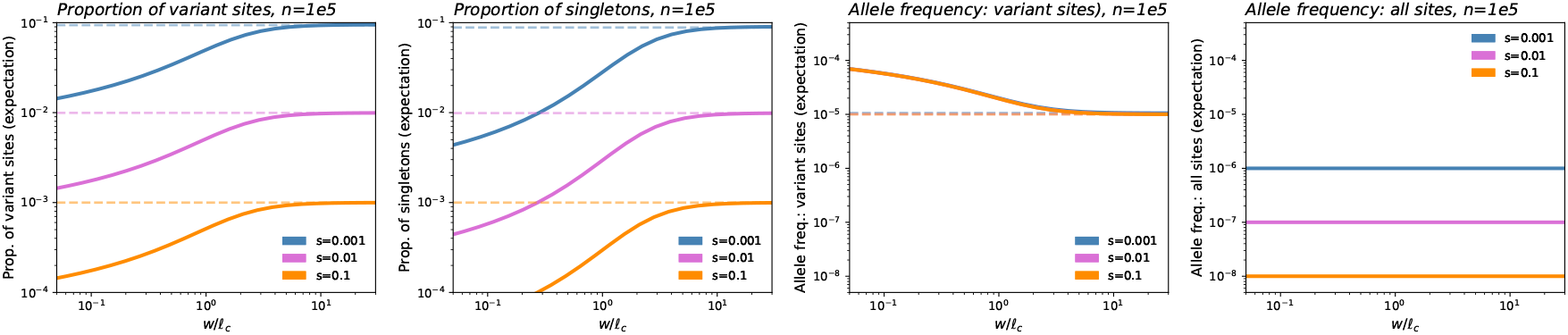
Summary statistics, as in Fig. 4, as a function the scaled sampling width *w*/𝓁_*c*_. Dashed lines represent theoretical expectation under uniform sampling. Inplots shown, *σ* = 10, *ρ*_*N*_ = 20, and *µ* = 1*e* − 9.

**Figure S12:**
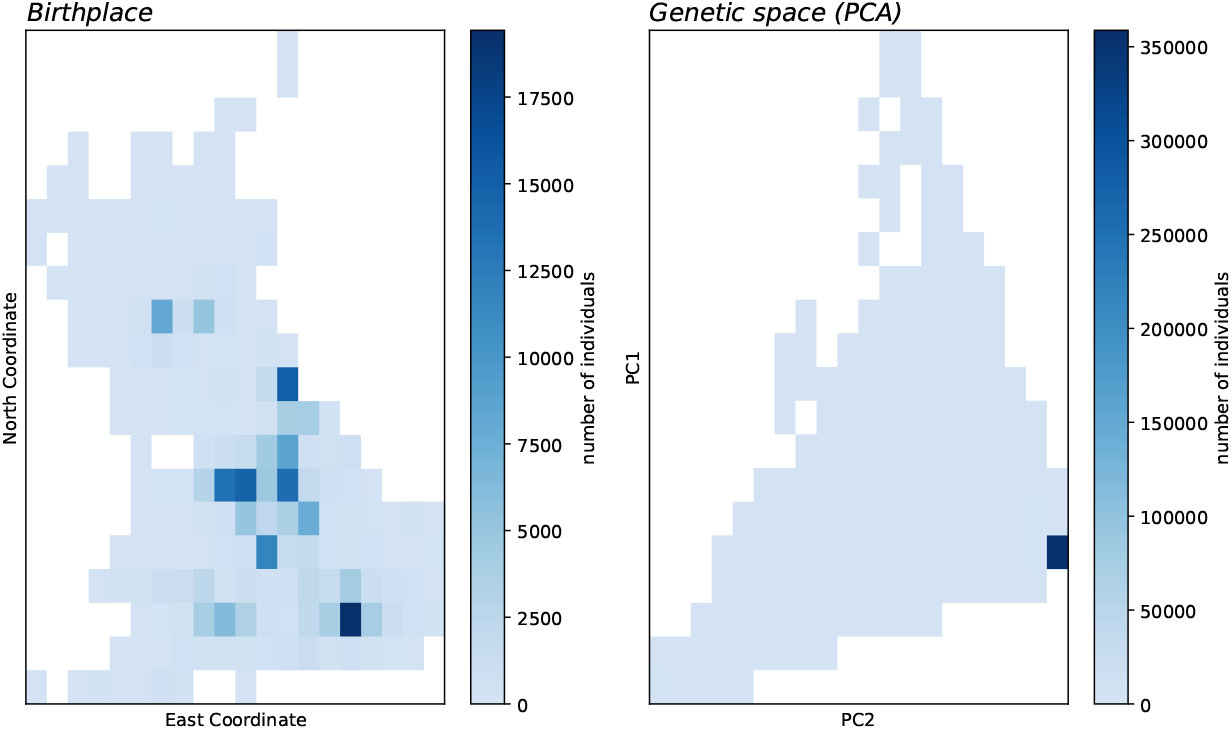
Counts of individuals in the UK Biobank included in empirical analyses over discretized geographic (left) and genetic (right) space. Each grid has dimensions 20×20 with equal-sized bins.

**Figure S13:**
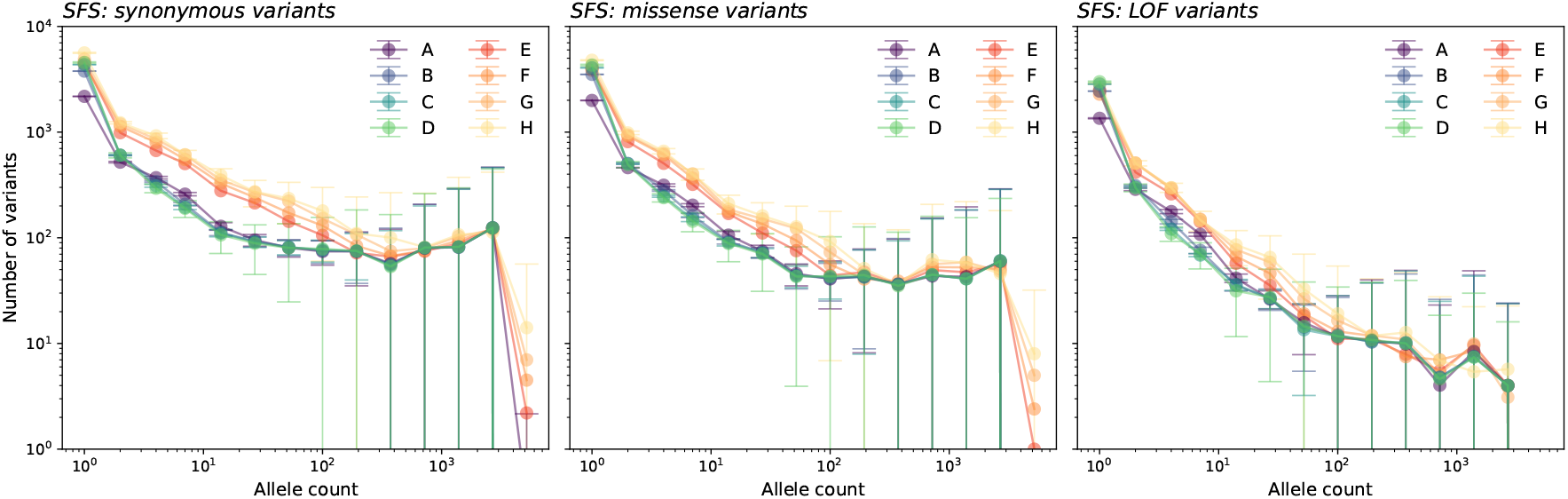
Site frequency spectra for sampling distributions as depicted in Fig. 3 across three variant classes.

**Figure S14:**
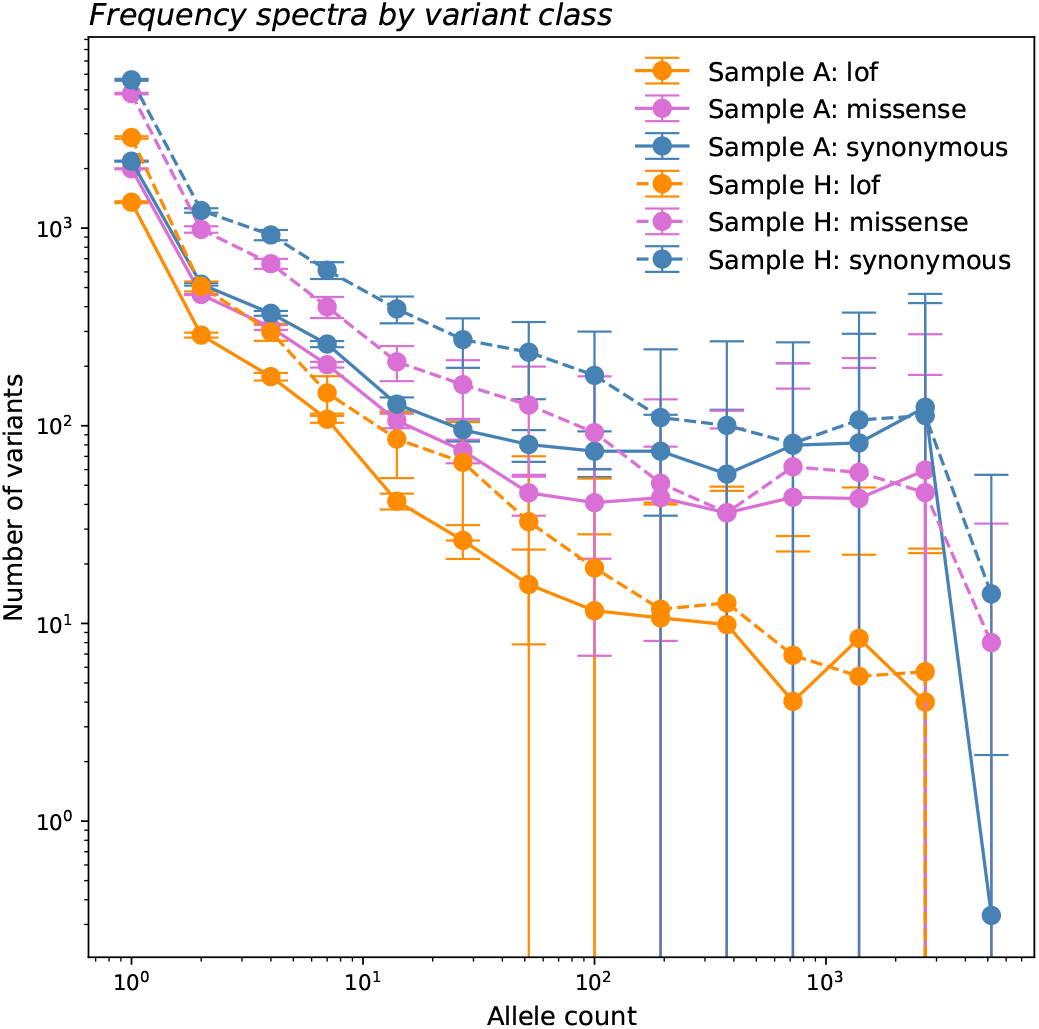
Site frequency spectra for narrowest (**A**) and broadest (**H**) sampling distributions across all three variant classes.

**Figure S15:**
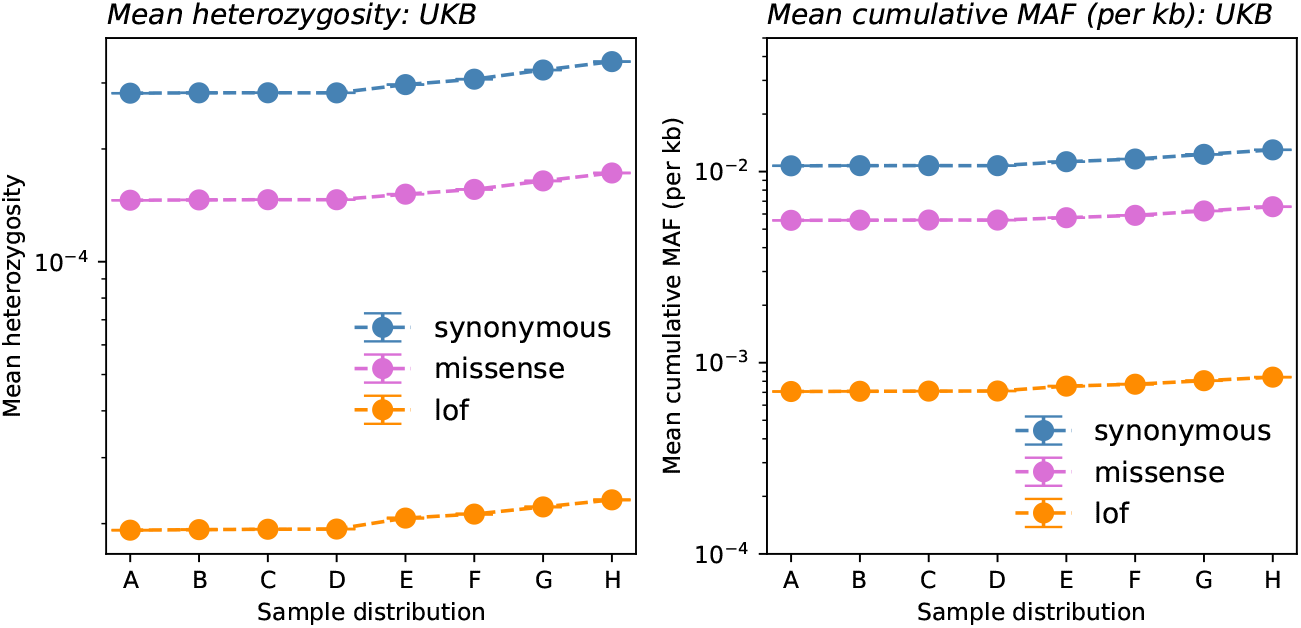
Mean heterozygosity (left) and cumulative MAF (per kb; right) as calculated from UKB re-samples.

**Figure S16:**
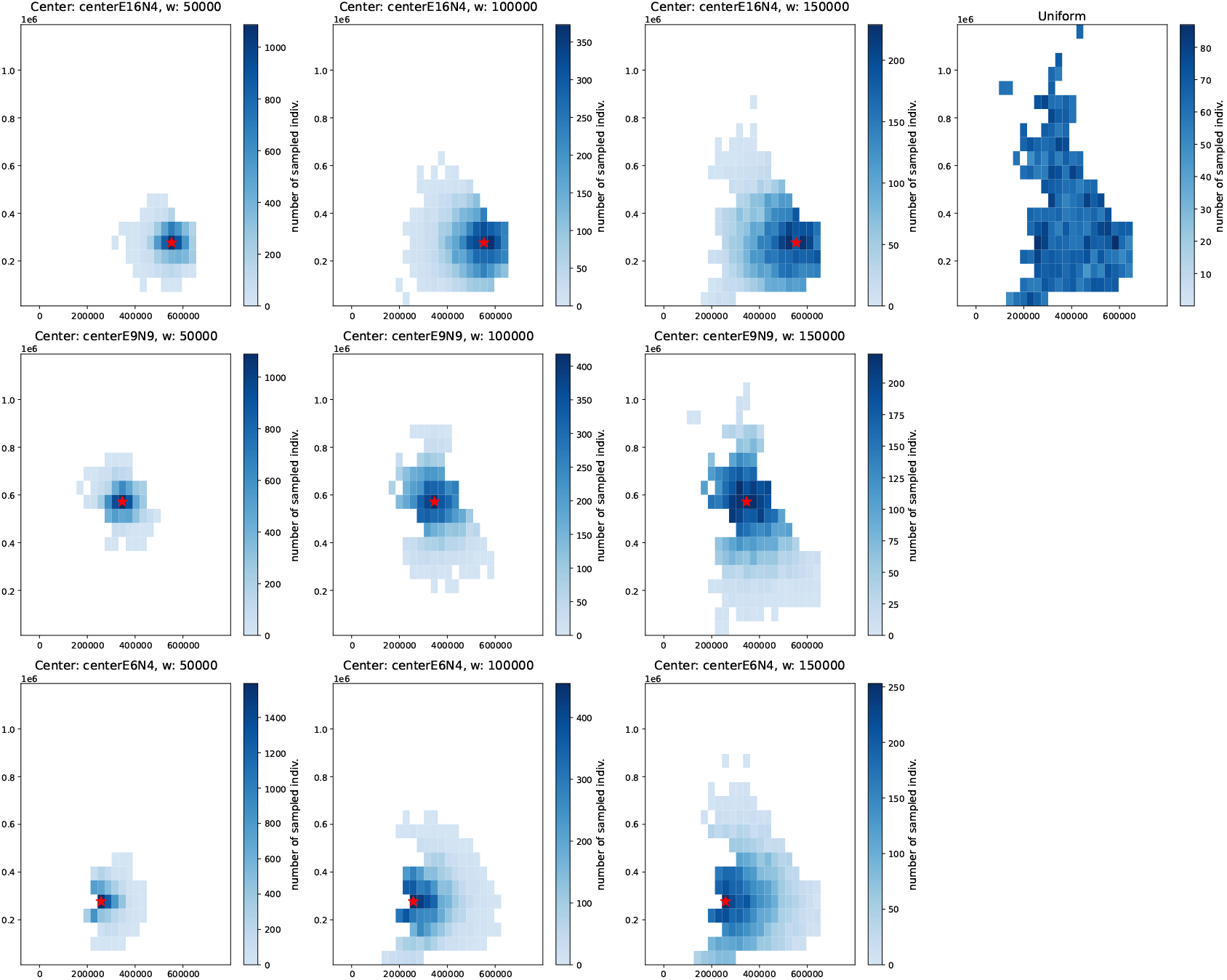
Visualization of sampling distributions in geographic space across three center points and values of *w* (as well as the uniform distribution).

